# IMPACTS OF DNA METHYLATION ON H2A.Z DEPOSITION AND NUCLEOSOME STABILITY

**DOI:** 10.1101/2025.07.31.667981

**Authors:** Rochelle M. Shih, Yasuhiro Arimura, Hide A. Konishi, Hironori Funabiki

## Abstract

The histone variant H2A.Z and DNA methylation are enriched at mutually exclusive genomic segments, though its mechanistic bases remain unclear. Here, we examine impacts of DNA methylation on the intrinsic stability of the H2A.Z nucleosome and chaperone-mediated H2A.Z deposition. Cryo-EM and endonuclease analyses suggest that DNA methylation subtly increases the openness and accessibility of the H2A.Z nucleosome on satellite II-derived DNA sequences. In transcriptionally silent *Xenopus* egg extracts, H2A.Z preferentially associates with unmethylated DNA though a substantial proportion of H2A.Z is recruited to methylated DNA. Preferential H2A.Z deposition to unmethylated DNA depends on the SRCAP complex, whose DNA binding is suppressed by methylation, while a SRCAP-independent and DNA methylation-insensitive mechanism for H2A.Z deposition also exists. Altogether, we propose that SRCAP drives the biased association of H2A.Z to unmethylated DNA, while additional mechanisms, potentially taking advantage of the subtle DNA methylation-induced physical effects, further assist the exclusion of H2A.Z from methylated DNA.

## INTRODUCTION

Structural features of the nucleosome, a protein-DNA complex composed of ∼146 bp DNA wrapped around a histone octamer, directly regulate genomic accessibility to a variety of DNA binding proteins, in part, through incorporation of histone variants with unique physical properties (Kurumizaka et al., 2021; Talbert and Henikoff, 2016). The highly evolutionarily conserved H2A variant, H2A.Z, is involved in transcriptional regulation with a well described localization at the transcription start sites (TSSs) of active genes (Bönisch and Hake, 2012; Giaimo et al., 2019; Zlatanova and Thakar, 2008). While typically associated with gene activation, H2A.Z has also been connected to a range of functions including DNA repair, chromosome segregation, heterochromatin maintenance, and transcriptional repression (Colino-Sanguino et al., 2022; Giaimo et al., 2019). Loss of H2A.Z induces lethality in many eukaryotes, such as mice, *Drosophila*, and *Tetrahymena* (Faast et al., 2001; Liu et al., 1996; Van Daal and Elgin, 1992), and dysregulation of its localization has been linked to several diseases, including various cancers and the developmental disorder Floating-Harbor syndrome (Buschbeck and Hake, 2017; Diegmüller et al., 2025). What mechanisms mediate proper H2A.Z localization, and how that connects with its broad-reaching functions, represents an important area of study.

Genomic sequencing analysis of a wide range of eukaryotes have found H2A.Z and DNA methylation, typically occurring as 5-methylcytosine (5mC), to be strikingly anti-correlated across the genome, at both a global and local level (Berta et al., 2021; Coleman-Derr and Zilberman, 2012; Conerly et al., 2010; Murphy et al., 2018; Nie et al., 2019; Zemach et al., 2010; Zilberman et al., 2008). The high conservation of this antagonism points to the existence of a shared fundamental pathway involved in creating this dichotomy and emphasizes its functional importance. In *Arabidopsis,* H2A.Z mediates recruitment of the demethylase ROS1, thereby promoting DNA demethylation; however, this pathway uses plant-specific machinery that targets only a particular set of genomic loci (Nie et al., 2019). More general mechanisms for how the mutually exclusive H2A.Z and DNA methylation compartments exist at large remain to be resolved.

DNA methylation may mediate H2A.Z exclusion via two distinct mechanisms: 1) influencing the intrinsic physical stability of the nucleosome, and/or 2) modulating the activity of nucleosome-remodelers. H2A.Z and H2A diverge in sequence along areas important for stabilizing intra-nucleosome interactions, such as the L1 loop, docking domain, and C-terminus (Bönisch and Hake, 2012). DNA methylation can influence the geometry of DNA, particularly its roll and twist, which then likely affect nucleosome wrapping (Li et al., 2022b). A large body of work has been conducted on the physical impacts of either H2A.Z or DNA methylation on the nucleosome, but conflicting conclusions have been reported; incorporation of H2A.Z or DNA methylation independently has been shown to cause stabilizing (Chen et al., 2013; Choy et al., 2010; Collings et al., 2013; Dai et al., 2021; Ishibashi et al., 2009; Lee et al., 2015; Lee and Lee, 2012; Park et al., 2004; Thambirajah et al., 2006), destabilizing (Horikoshi et al., 2019; Jimenez-Useche et al., 2014; Jimenez-Useche and Yuan, 2012; Lee et al., 2015; Lewis et al., 2021; Li et al., 2023; Ngo et al., 2016; Rudnizky et al., 2016; Weber et al., 2010; Zhang et al., 2005), or no detectable effects (Fujii et al., 2016; Langecker et al., 2015; Osakabe et al., 2015). Nevertheless, it is possible that the combination of H2A.Z and DNA methylation’s effects may physically bias against retention of H2A.Z nucleosomes on methylated DNA and contribute to their overall antagonism.

H2A.Z remodeling is primarily carried out by the Ino80 family of nucleosome remodelers. Both the SRCAP complex (SRCAP-C) and TIP60 complex (TIP60-C) mediate exchange of canonical H2A to H2A.Z within a nucleosome (Ruhl et al., 2006), while the INO80 complex (INO80-C) has been reported to evict H2A.Z (Scacchetti and Becker, 2021; Watanabe and Peterson, 2010). TIP60-C can also associate with the H2A.Z chaperone, ANP32e, to evict H2A.Z from nucleosomes (Mao et al., 2014; Obri et al., 2014). Members of this remodeler family are prime candidates for enforcing the separation of H2A.Z and DNA methylation; however, it is unknown whether any of these complexes exhibit DNA methylation sensitivity.

To determine molecular mechanisms contributing to the mutual exclusion of H2A.Z and DNA methylation within the genome, here we investigated the effects of DNA methylation on both the physical stability of H2A.Z nucleosomes and remodeler-mediated deposition of H2A.Z onto synthetic DNA constructs by employing cryo-EM structural analysis and the physiological *Xenopus* egg extract system, respectively.

## RESULTS

### Cryo-EM analysis of H2A.Z nucleosomes in the presence or absence of DNA methylation

To assess the physical impacts of DNA methylation on H2A.Z nucleosomes, we solved cryo-EM structures of human H2A.Z nucleosomes reconstituted with either methylated (Me) or unmethylated (UM) Sat2R-P DNA (**Fig 1, Suppl Fig 1, Suppl Table 1**). The Sat2R-P sequence is a 152 bp artificial palindromic sequence containing 9 evenly distributed CpG sites and was engineered from a published canonical nucleosome crystal structure containing a methylated derivative of the human satellite II sequence (termed Sat2R) (Osakabe et al., 2015) (**Fig 1A**). This sequence was chosen for three reasons: 1) human satellite II (HSat2) represents a highly abundant methylated sequence which undergoes disease-associated differential methylation (Ehrlich, 2003; Jackson et al., 2004; Tilman et al., 2012; Unoki, 2021; Wong et al., 2001), 2) it was reported that little H2A.Z associates with HSat2 repeats (Capurso et al., 2012), which are frequently hypomethylated in cancers (Ehrlich, 2009; Hall et al., 2017), and 3) its symmetric palindromic sequence was needed to increase final structure resolutions due to the two-fold pseudo symmetry about the nucleosome dyad axis. Prepared nucleosomes were run on a native PAGE gel to assess for quality, where both samples appeared as a singular band with minor comparable populations of unincorporated DNA (**Suppl Fig 1A**).

**Figure 1.**
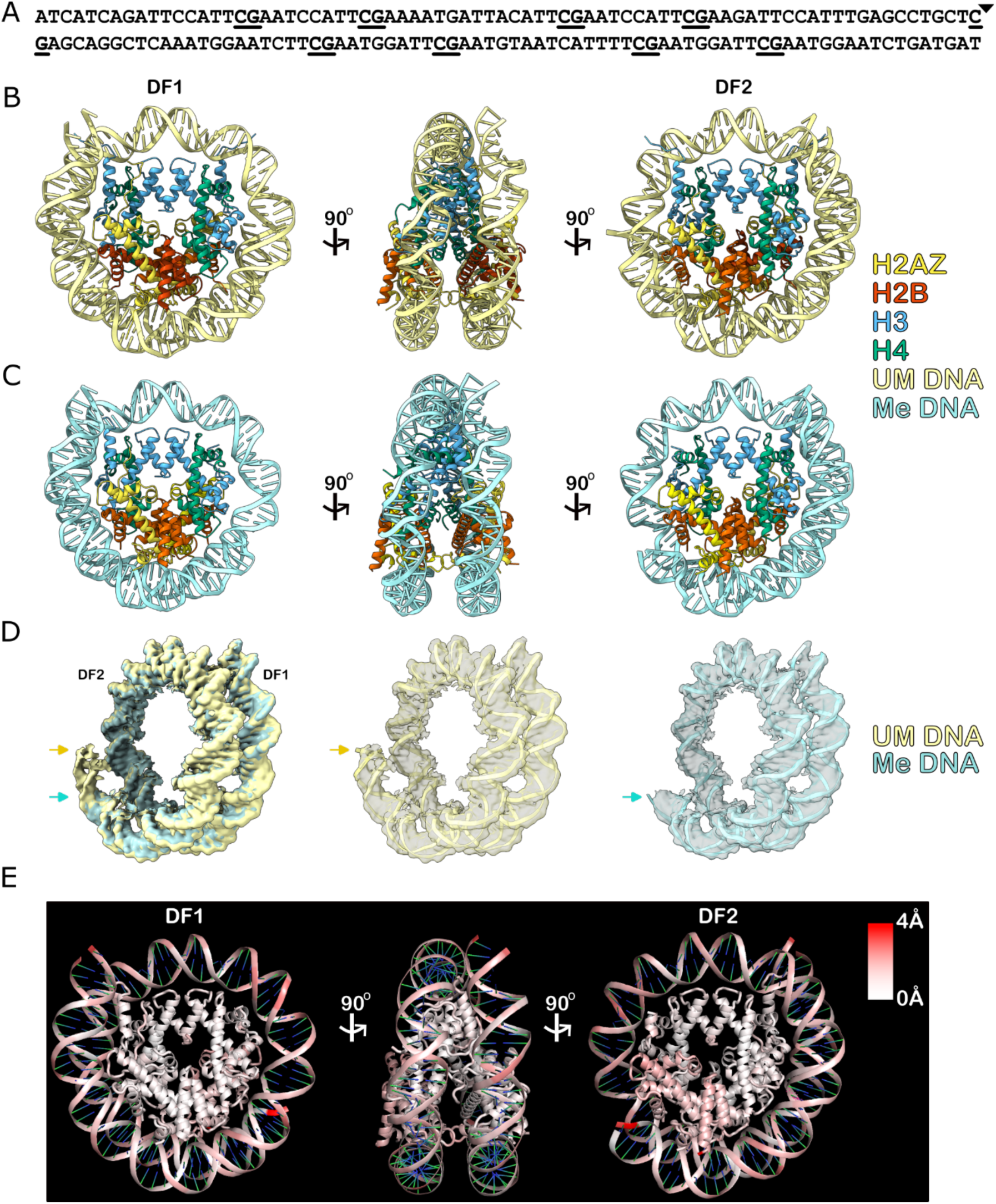
Cryo-EM structures of methylated and unmethylated Sat2R-P human H2A.Z nucleosomes. **(A)** Diagram of the palindromic 152 bp Sat2R-P sequence. CpGs are underlined. Triangle denotes midpoint. **(B and C)** Atomic models of unmethylated (**B**) and methylated (**C**) Sat2R-P H2A.Z structures (the v1 model for both). Two face views (DF1 and DF2) and a side view (middle) are shown. **(D)** Left, overlay of unmethylated (UM) and methylated (Me) Sat2R-P DNA EM densities. Middle and right, overlays of DNA densities on top of the corresponding v1 DNA atomic models for either the unmethylated (middle) or methylated (right) structures. Arrows point to the visualizable end of the DF2 linker DNA for each structure. **(E)** RMSD analysis comparing differences between unmethylated and methylated v1 Sat2R-P H2A.Z atomic models.

Cryo-EM structures of UM and Me Sat2R-P H2A.Z nucleosomes were solved at a final resolution of 3.05 Å and 2.78 Å, respectively (**Fig 1B and C, Suppl Fig 1**). We found that the nucleosome dyad was shifted roughly two base pairs from the center of the Sat2R-P palindrome, introducing a sequence asymmetry despite our intended use of the palindromic sequence. The shifted nucleotide positions were determined using the higher resolution densities surrounding the nucleosome dyad, which allowed for pyrimidine and purine distinction (**Suppl Fig 2A-D**). Two atomic models (termed v1 and v2) with mirrored DNA sequences were generated for each structure to account for the resultant sequence asymmetry, with the final EM densities likely reflecting an average of these two models (**Suppl Fig 2E**).

As a whole, both the methylated and unmethylated structures resembled previously published H2A.Z nucleosome structures and exhibited a linker DNA asymmetry where one side of the nucleosome (distal face 2, DF2) contained a highly flexible linker DNA region compared to the other (distal face 1, DF1) (**Fig 1B and C**) (Horikoshi et al., 2013; Lewis et al., 2021; Suto et al., 2000). As a palindromic DNA sequence was used, it is likely that the asymmetrical faces in the cryo-EM structures arose due to an averaging of closed and open H2A.Z nucleosome particles into the same structure rather than representing an asymmetry found in individual particles.

In a direct comparison of DF2 linker DNA lengths, we found that the Me Sat2R-P map had much weaker linker DNA density than UM Sat2R-P, resulting in a ∼7 bp shorter resolved DNA model for the methylated structure (**Fig 1D**). This loss of linker density indicates destabilized histone-DNA interactions and increased DNA structural variability in the presence of DNA methylation. Along similar lines, Root Mean Squared Distance (RMSD) analysis identified the H2A.Z-H2B dimer of DF2, as well as the overall DNA backbone, as regions of considerable difference between the methylated and unmethylated structures, whereas DF1, the closed side of the nucleosome with a highly resolved linker DNA, was largely similar between the two (**Fig 1E**).

### DNA methylation creates open H2AZ nucleosome structures

To further characterize areas of change, maps of unmethylated and methylated H2A.Z structures were filtered and resampled to the same resolution (∼3 Å) for fair comparison (**Fig 2A**). Three major differences between the maps were identified. First, cryo-EM densities of the H3 N-terminus and H2A.Z C-terminus on DF2 were much weaker in the methylated structure as compared to unmethylated, preventing model construction in these areas (**Fig 2B**). Second, an extra density on the H2A.Z-H2B acidic patch was seen on the methylated map but was not present on the unmethylated map (**Fig 2C**). Third, between the two major orientations the H4 N-terminal tail can adopt (Arimura et al., 2021), the unmethylated map showed a strong preference for the inward orientation whereas the methylated structure exhibited similar populations of inward and outward orientations (**Fig 2D**).

**Figure 2.**
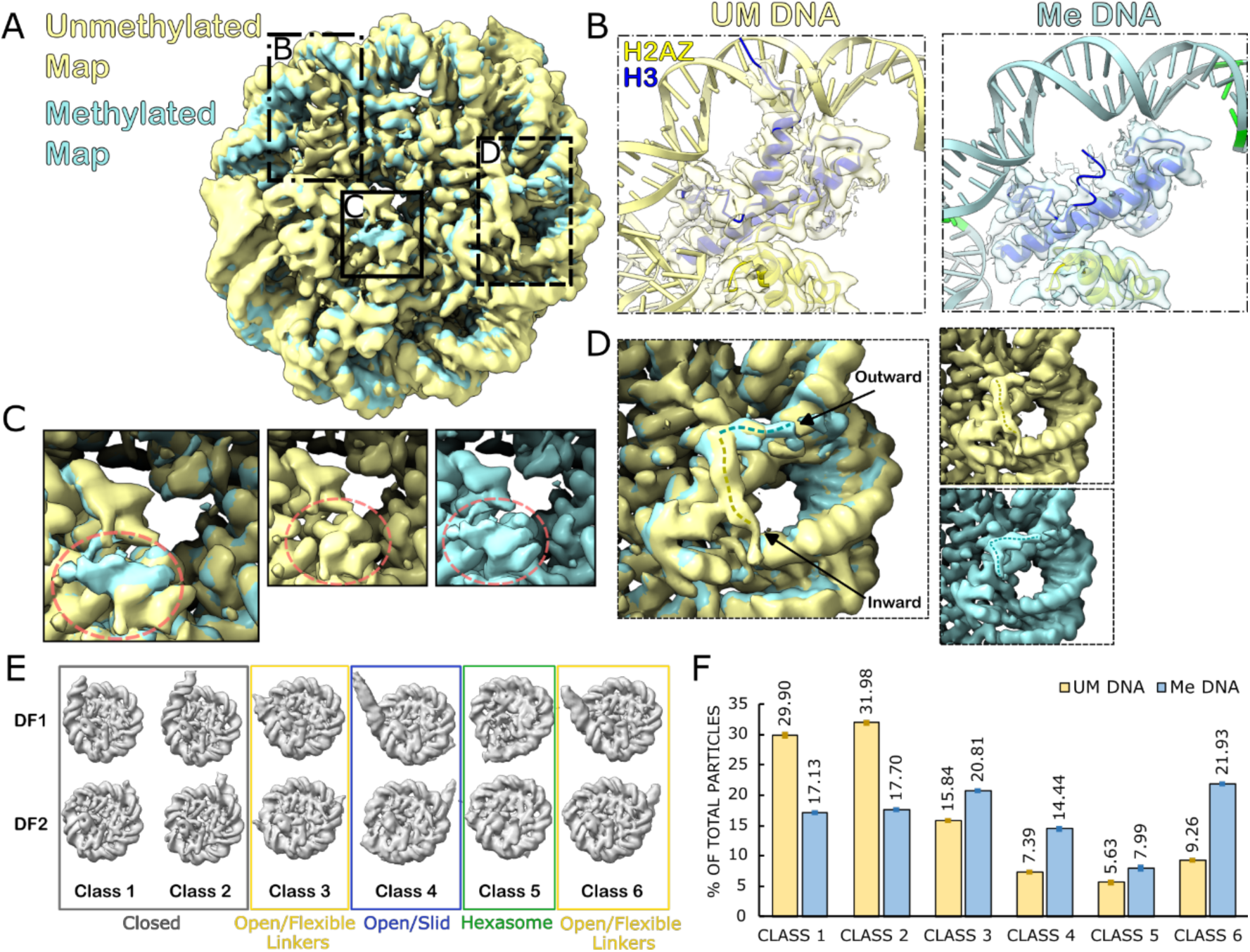
DNA methylation reduces H3-DNA contacts, alters H4 N-terminal tail orientations, and forms open nucleosome structures. **(A)** Density maps of unmethylated (UM) and methylated (Me) Sat2R-P H2A.Z nucleosomes filtered and resampled to ∼3Å. The face shown, and all following zoom-ins, is of DF2. Maps are overlaid onto each other. Boxed regions highlight major areas of difference shown in **B-D**. **(B)** Zoom-in of the H3 N-terminus on DF2 of both structures. Density maps are shown as a transparent overlay on top of the corresponding v1 atomic models. **(C)** Zoom-in of the DF2 acidic patch on both structures. Area corresponding to the extra density found in the Me structure labeled in red. **(D)** Zoom-in of the H4 N-terminal tail. Dotted lines show each structure’s preferred orientation **(E)** 3D Models used in the *in silico* mixing 3D classification analysis. Classes are as follows: (1 and 2) closed nucleosome structures with varying linker lengths; (3 and 6) open nucleosomes with highly flexible linkers; (4) open nucleosomes that have shifted position from center; (5) hexasomes. Both distal faces (DF1 and DF2) are shown of each model. **(F)** Results of classification analysis. Y-axis is the proportion of particles sorted into each class as a percent of the total input particles from each sample (UM or Me). Error bars represent SEM, n = 3 technical replicates.

The H3 N-terminus interacts with the H2A/H2A.Z C-terminus and DNA to hold the linker DNA close to the body of the nucleosome (Li et al., 2023; Li and Kono, 2016). Loss of the H3 N-terminal density from the methylated structure suggests destabilized DNA-histone interactions that are consistent with the shorter resolved linker DNA on DF2 of the methylated model compared to unmethylated (**Fig 1D**). The H4 N-terminal tail, on the other hand, mediates inter-nucleosomal interactions through contacts with a neighboring acidic patch (Chen et al., 2017; Luger et al., 1997). The extra density on the Me Sat2R-P acidic patch may then have come from the H4 tail of another nucleosome, in line with the changes also observed in H4 tail orientation. These observations may be consistent with reports that DNA methylation increases condensation of nucleosomes, as well as bare DNA, *in vitro* and *in vivo* (Buitrago et al., 2021; Jimenez-Useche et al., 2014; Yang et al., 2020), though require future experimental validation.

To more quantitatively assess methylation-dependent structural changes, we conducted an *in silico* mixing 3D classification analysis (Arimura et al., 2021; Hite and MacKinnon, 2017), by combining micrographs collected from both methylated and unmethylated nucleosome samples and then sorted particles picked from the combined micrographs into six 3D classes of nucleosomes, each with varying degrees of open or closed linker DNA (**Fig 2E and F, Suppl Fig 3**). Most of the unmethylated particles (>60%) were sorted into the closed linker classes (Class 1 and 2) whereas the majority of methylated particles (∼60%) belonged to classes with open linkers (Classes 3, 4, 6) (**Fig 2F**). The fraction of hexasomes (Class 5) is minor in both conditions with a slight increase in methylated particles (Class 5). Among these open linker classes that are preferentially found in methylated particles, Class 4 also represents nucleosomes where the palindromic center of the DNA has slid and is considerably offset from the dyad axis of the histone octamer. This may explain the weaker DF2 linker density seen in the high resolution Me Sat2R-P structure (**Fig 1D**). Additionally, the DNA backbone of the methylated nucleosome structure had a lower local resolution compared to unmethylated, despite the methylated map having a higher global resolution (**Suppl Fig 4**). This is consistent with the increased population of open linker and slid nucleosome particles in the methylated structure. Altogether, our cryo-EM structural analysis suggests that DNA methylation creates open H2A.Z nucleosome structures with destabilized DNA wrapping.

### Cryo-EM structures of 601L H2A.Z nucleosomes show no difference between methylation status

In addition to the Sat2R-P nucleosomes, we also solved cryo-EM structures of human H2A.Z nucleosomes reconstituted on methylated and unmethylated 601L DNA, a 205 bp palindromic version of the lefthand portion of the classic Widom 601 sequence (**Suppl Fig 5 and 6, Suppl Table 1**) (Lowary and Widom, 1998). Final maps of the unmethylated and methylated structures were obtained at 3.09 Å and 3.25 Å, respectively (**Suppl Fig 5B-E**). RMSD analysis of the solved structures showed that both maps were essentially identical to each other, with the few differences present localized to highly flexible regions of the nucleosome (**Suppl Fig 5F**). The lack of clear structural differences when using 601L DNA is likely due to its artificially strong nucleosome positioning ability, which may have prevented detection of subtle changes caused by DNA methylation.

### DNA methylation increases accessibility of H2A.Z but not H2A nucleosomes

In order to validate the observed opening of the H2A.Z nucleosome structure by DNA methylation, and to determine if this methylation effect is also present on nucleosomes containing canonical H2A, we measured nucleosome DNA accessibility via restriction enzyme digest (**Fig 3, Suppl Fig 7 and 8**). We tested both human H2A and H2A.Z nucleosomes reconstituted with methylated and unmethylated versions of 152 bp non-palindromic Sat2R DNA that was modified to include a singular HinfI site 18 bp in from the 5’-end containing a Cy5-fluorophore (**Fig 3A**). Based on our Sat2R-P H2A.Z nucleosome cryo-EM structures as well as published Sat2R H2A nucleosome crystal structures(Osakabe et al., 2015), this HinfI site is predicted to be located at SHL5.5, where the L2 loop of H2A/H2A.Z contacts the DNA minor groove (**Fig 3B**). Since this SHL5.5 was only resolved in DF1 but not DF2 of the methylated Sat2R-P H2A.Z structure, whereas it was resolved in both DF1 and DF2 in UM Sat2R-P H2A.Z nucleosomes, we assumed that this segment would show methylation-induced accessibility changes to the HinfI endonuclease.

**Figure 3.**
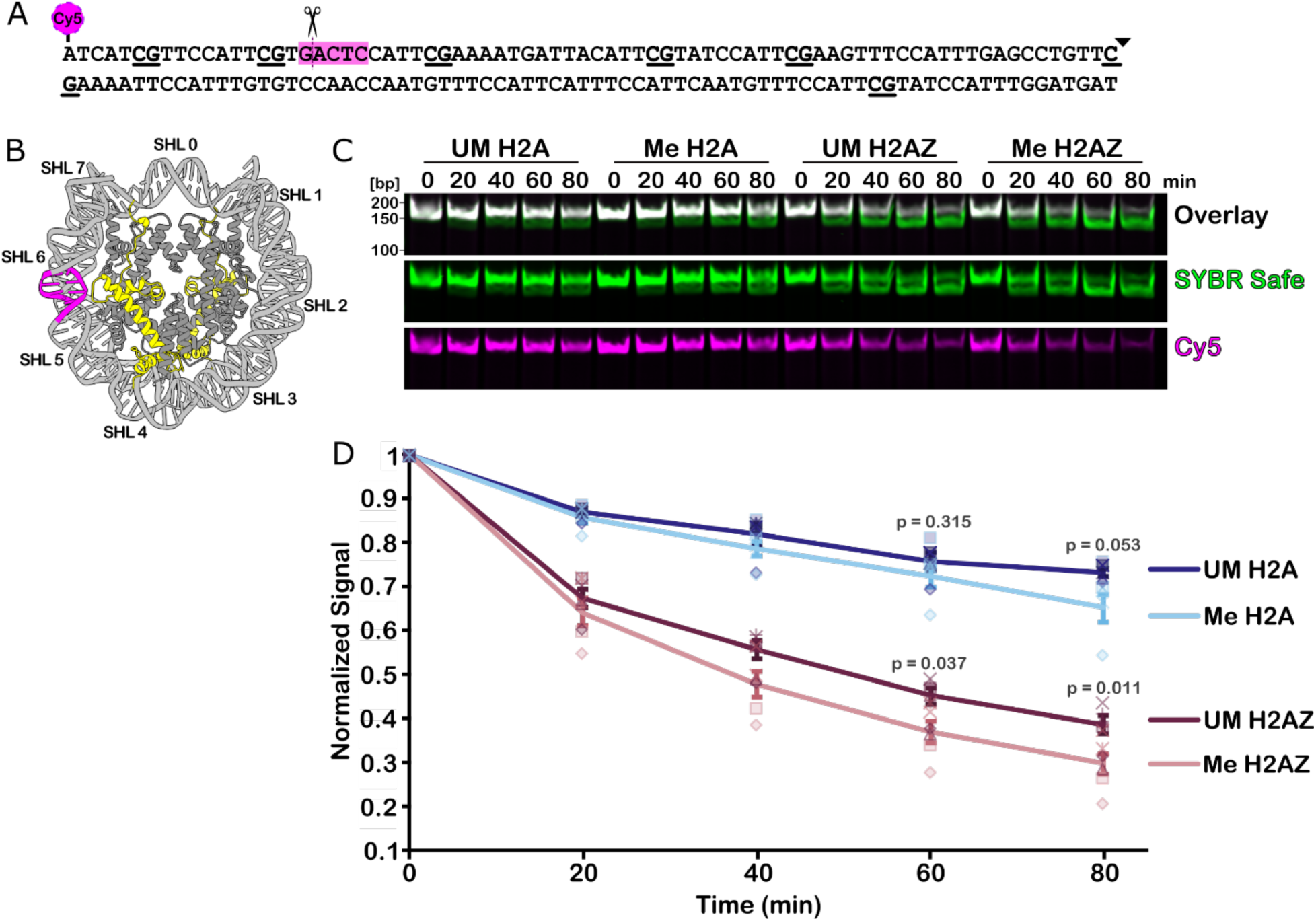
DNA methylation increases H2A.Z nucleosome accessibility. **(A)** Diagram of the **15**2 bp 1HinfI_Sat2R DNA sequence used. CpG sites underlined and sequence midpoint indicated by the triangle. HinfI recognition site highlighted in magenta. A Cy5 fluorophore is attached to the 5’ end nearest the HinfI site. **(B)** Schematic of 1HinfI_Sat2R H2A.Z nucleosome with HinfI recognition site demarcated in magenta and H2A.Z in yellow. **(C)** Native gel showing HinfI digestion time course of UM and Me human H2A and H2A.Z nucleosomes. SYBR Safe was used to stain total DNA. Digestion efficiency was assessed via loss of Cy5 signal and a downward shift in the total DNA band. **(D)** Quantification of (**C**). Cy5 signals were normalized to total SYBR Safe signal for each sample. Error bars represent SEM. Each shape represents data from one independent experiment, n = 5 experiments. Statistics comparing unmethylated and methylated samples at indicated timepoints were completed with a two-tailed t-test assuming unequal variance with *p*-values listed above each comparison.

Homogeneity of the reconstituted nucleosomes was assessed via native PAGE and mass photometry analysis prior to restriction digest analysis (**Suppl Fig 7A-C**). Digest of bare control DNA confirmed that HinfI behaved similarly on both methylated and unmethylated Sat2R substrates over the time scale used in the experiment (**Suppl Fig 7D**). We found that H2A.Z nucleosomes were more accessible to enzymatic digest than H2A nucleosomes, independent of DNA methylation status (**Fig 3B and C, Suppl Fig 8**). This result was consistent with previous reports using similar assays on Widom 601 nucleosomes, showing that the DNA termini of H2A.Z mono-nucleosomes are more accessible than that of H2A (Jung et al., 2024; Lewis et al., 2021). When comparing methylation-dependent differences for each type of nucleosome, methylated and unmethylated H2A nucleosomes were largely digested at similar rates. Methylated H2A.Z nucleosomes, on the other hand, exhibited a small but more pronounced increase in accessibility as compared to its unmethylated counterpart (**Fig 3B and C, Suppl Fig 8**). These results corroborate our structural findings and suggest that the effects of DNA methylation may be negligible on an intrinsically stable nucleosome (i.e. containing H2A or with hyper-stable nucleosome positioning sequences) but can enhance nucleosome opening on an already open and accessible H2A.Z nucleosome.

### H2A.Z and DNA methylation localization in *Xenopus laevis* sperm pronuclei and a fibroblast cell line

Results from our structural and biochemical experiments indicate that DNA methylation subtly increases the accessibility of H2A.Z nucleosomes. While functional implications likely exist, this slight effect on the intrinsic physical stability of the H2A.Z nucleosome is unlikely to be the sole driver of the mutually exclusive H2A.Z and DNA methylation compartmentalization in cells. To identify the major biochemical mechanism underlying this dichotomy, we turned to the *Xenopus laevis* egg extract system, a unique cell-free system that allows for characterization of chromatin proteins under physiological conditions (Almouzni and Wolffe, 1993; Hoogenboom et al., 2017; Jenness et al., 2018; Lohka and Maller, 1985; Onikubo and Shechter, 2016; Postow et al., 2008; Wassing et al., 2024; Xue et al., 2013; Zierhut et al., 2014).

To ensure that *Xenopus* egg extract is a valid tool for studying the H2A.Z-DNA methylation antagonism, we conducted CUT&Tag – Bisulfite Sequencing (CnT-BS) (Li et al., 2021), a method which allows for the direct assessment of DNA methylation on CUT&Tag reads, against H2A.Z of *Xenopus* sperm nuclei incubated in interphase extract. This condition mimics fertilization and allows for full nucleosome assembly onto sperm (primarily composed of protamines and H3/H4 tetramers) to create functional pronuclei (Oikawa et al., 2020). We also collected samples against H3 to serve as a pan-nucleosome control. Since sperm pronuclei in egg extract represents a unique transcriptionally silent state (Engelke et al., 1980; Newport and Kirschner, 1982; Veenstra et al., 1999), we also performed CnT-BS on XTC-2 cells, a *Xenopus laevis* fibroblast cell line (Pudney et al., 1973), to assess changes in DNA methylation and H2A.Z overlap across different nuclei types.

Initial comparisons of CnT-BS reads across two biological replicates of our four conditions demonstrated good reproducibility between replicates, while each set of experimental conditions exhibited distinct characteristics. A greater than 99 % bisulfite conversion efficiency rate, as assessed through our unmethylated lambda DNA spike-in, was achieved, and correlation matrices of both the mapped reads and methylation calls were highly similar between biological replicates (**Suppl Fig 9, Suppl Table 2 and 3**). When comparing across conditions, for both the H3/H2A.Z mapped reads and DNA methylation sites, the first degree of difference seen was between XTC-2 and sperm samples, and then H3 vs H2A.Z following after, indicating that large-scale changes in the DNA methylation and genomic landscape occurred between the two different nuclear states (**Suppl Fig 9**).

To assess characteristics of H2A.Z and H3 genomic distributions, we subset the *Xenopus* genome into 1,000 bp bins and classified the top 20 % of bins containing the most mapped reads for each condition based on annotated functionalities (Fisher et al., 2023). We found that a greater proportion of highly mapped H2A.Z bins were located near promoter regions as compared to H3 for both sperm and XTC-2 samples (**Suppl Fig 10A**). Although transcription is suppressed in *Xenopus* egg extracts, H2A.Z in sperm pronuclei were well positioned at TSSs, as generally seen for actively transcribed genes in somatic cells (**Suppl Fig 10B and C**). This mirrors the reported deposition of H3K4me3 at the +1 TSS sites in *Xenopus* sperm nuclei prior to remodeling as well as reports in zebrafish sperm and *Drosophila* that H2A.Z is positioned at promoters prior to zygotic genome activation (ZGA) (Ibarra-Morales et al., 2021; Murphy et al., 2018; Oikawa et al., 2020).

### H2A.Z is biased towards hypomethylated regions of the genome in both *Xenopus* sperm pronuclei and the XTC-2 fibroblast cell line

We next determined the methylation status of CpGs associated with either H2A.Z or H3 across replicates of XTC-2 and sperm nuclei (**Fig 4A**). One notable drawback of CUT&Tag is a bias towards open and accessible regions (Kaya-Okur et al., 2020; Richard et al., 2025), which may artificially amplify open chromatin signals and lead to an underestimation of DNA methylation status. To circumvent this issue, we analyzed H3 CpG methylation in areas outside of H2A.Z peaks (which tend to be highly accessible) to compare genomic methylation profiles in H2A.Z low/absent regions (H3) vs H2A.Z enriched peaks (H2A.Z).

**Figure 4.**
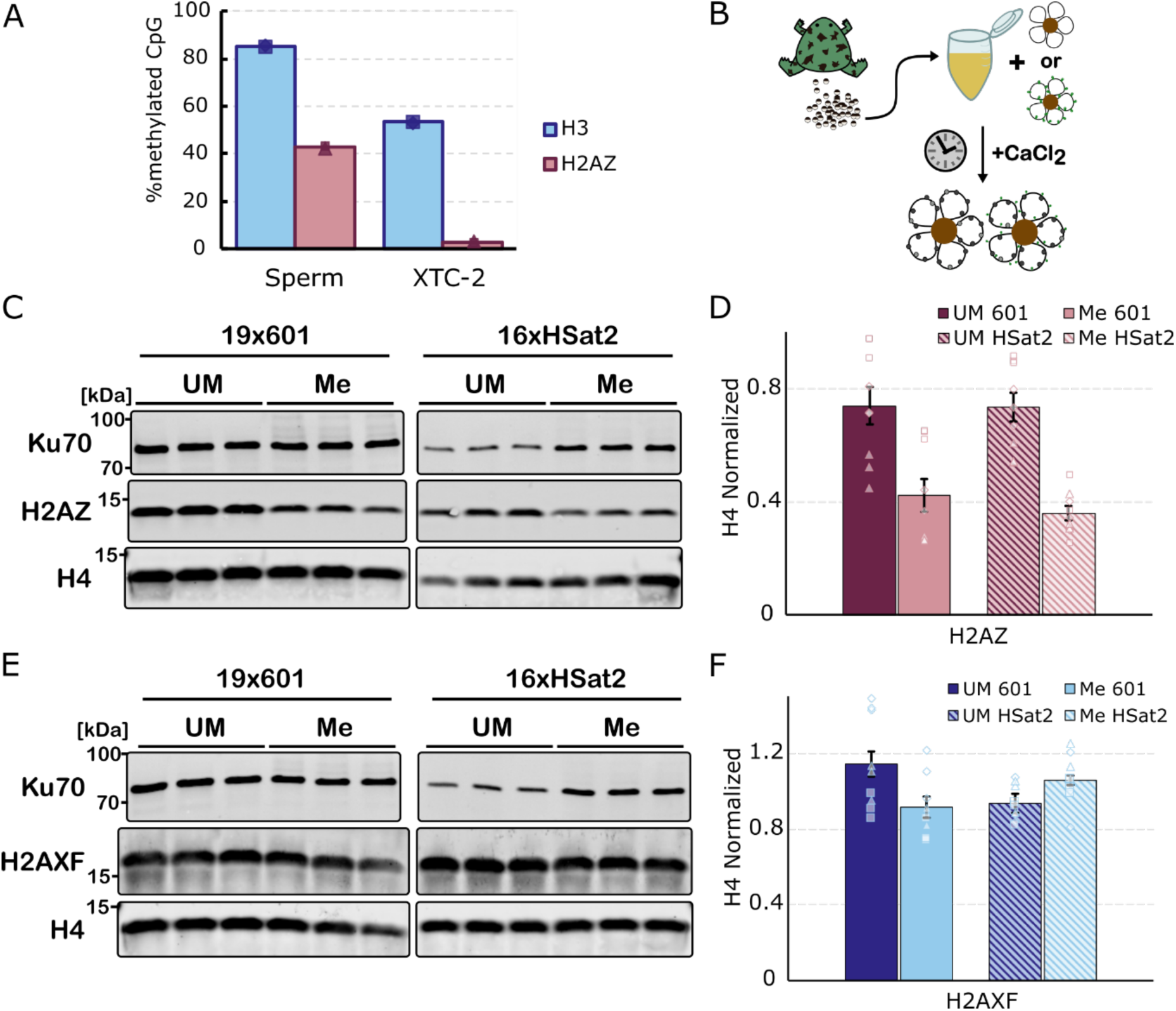
Presence or absence of DNA methylation influences H2A.Z deposition. **(A)** Percentage of methylated CpG sites (≥ 5x coverage) associated with H2A.Z peaks or H3 reads in sperm pronuclei incubated in interphase egg extract or XTC-2 nuclei following CnT-BS library preparation. H3 data is taken from regions outside of H2A.Z enriched peaks. Data points represent biological replicates (n = 2). **(B)** Schematic of chromatinization assay. Magnetic beads coated with DNA of interest were incubated in *Xenopus* egg extract with the addition of CaCl_2_ to induce cycling into interphase. After 60 min, DNA beads were isolated to assess for histone composition. **(C and E)** Western blot results of chromatinization assay probing for either H2A.Z (**C**) or H2A.X-F (**E**). Ku70 and H4 signals are shown as loading controls. Beads coated with unmethylated (UM) or methylated (Me) 19-mer arrays of Widom 601 (19×601) or 16-mer arrays of HSat2 (16xHSat2) were used. A representative blot of technical triplicates for each condition is presented. **(D and F)** Quantification of (**C** and **E**). H2A.Z and H2A.X-F signals were normalized to H4 intensity. Error bars represent SEM. Data points represent technical triplicates across 3 biological replicates with each shape representing data from a single independent experiment (n = 9).

Overall, sperm chromatin had a higher degree of methylation than XTC-2 chromatin, with 85.3 ± 0.4 % *s.d.* of CpGs in H3 reads experiencing methylation in sperm compared to 53.5 ± 0.4 % in XTC nuclei (**Suppl Table 2**). Methylated CpG frequencies of H2A.Z-associated reads were reduced to 43.0 ± 0.7 % in sperm chromatin and even further down to 3.0 ± 0.3 % in XTC-2 cells. Mapping of H3 associated methylation calls to annotated genes of the *Xenopus laevis* genome showed ubiquitous distribution of methylation across the length of genes, except for at the TSSs where methylation was depleted (**Suppl Fig 10B and C left**). Nearly all of these H3-marked hypomethylated TSSs exhibited H2A.Z enrichment in both sperm pronuclei and XTC-2 (**Suppl Fig 10B and C right**), indicating that hypomethylation-exclusive positioning of H2A.Z is most prominent at the TSSs (**Suppl Fig 10D**).

### DNA methylation-sensitive H2A.Z deposition on synthetic DNA constructs in *Xenopus* egg extract

To directly test whether the presence or absence of DNA methylation can dictate preferential H2A.Z deposition independently of other pre-existing epigenetic marks or DNA sequence, we took advantage of the *Xenopus* egg extract’s ability to efficiently assemble nucleosomes *de novo* onto naked DNA templates (Laskey et al., 1977) (**Fig 4B**). We found that, upon incubation of DNA coated magnetic beads in interphase egg extract, H2A.Z accumulated less efficiently on methylated DNA substrates compared to unmethylated (**Fig 4C and D, Suppl Fig 11**), whereas this methylation-sensitive deposition was not seen for H2A.X-F, the predominant H2A histone variant in *Xenopus* egg extract (**Fig 4E and F, Suppl Fig 11**) (Shechter et al., 2009). The methylation bias of H2A.Z held true for arrays of either the GC-rich Widom 601 (57 % GC content) or the more AT-rich human HSat2 (38% GC content) sequence. We next asked whether both H2A.Z variants, H2A.Z.1 and H2A.Z.2 (H2A.V), exhibit the same DNA methylation sensitivity. Since our antibody does not distinguish between the two H2A.Z paralogs, we translated ALFA-tagged (Götzke et al., 2019) versions of the variants in egg extract from exogenously added mRNA and found that both paralogs displayed preferential deposition onto unmethylated DNA (**Suppl Fig 12**).

### The SRCAP complex mediates H2A.Z’s preferential deposition on unmethylated DNA

Since SRCAP-C is known to play a major role in depositing H2A.Z to nucleosomes via exchange of H2A dimers on the nucleosome with H2A.Z (Ruhl et al., 2006), we examined if the observed preferential association of H2A.Z with unmethylated DNA in *Xenopus* egg extracts is mediated by SRCAP-C. Depletion of SRCAP-C from extract using a custom antibody against *Xenopus laevis* SRCAP greatly reduced the amount of H2A.Z loaded onto unmethylated DNA compared to control depletion (ΔIgG), while H2A.Z levels on methylated DNA were not affected by SRCAP depletion (**Fig 5A-C, Suppl Fig 13A-C**). Consequently, in SRCAP-C-depleted (ΔSRCAP) extracts, residual H2A.Z binding to DNA became insensitive to DNA methylation status (**Fig 5B and C, Suppl Fig 13C**). The SRCAP depletion effect was specific to H2A.Z since the amount of H2A.X-F, the predominant H2A variant in *Xenopus* eggs (Shechter et al., 2009), on DNA was not affected by SRCAP-C depletion (**Fig 5B and D, Suppl Fig 13C**). Similar results were reproduced with use of a commercially available antibody raised against human SRCAP (**Suppl Fig 14**).

**Figure 5.**
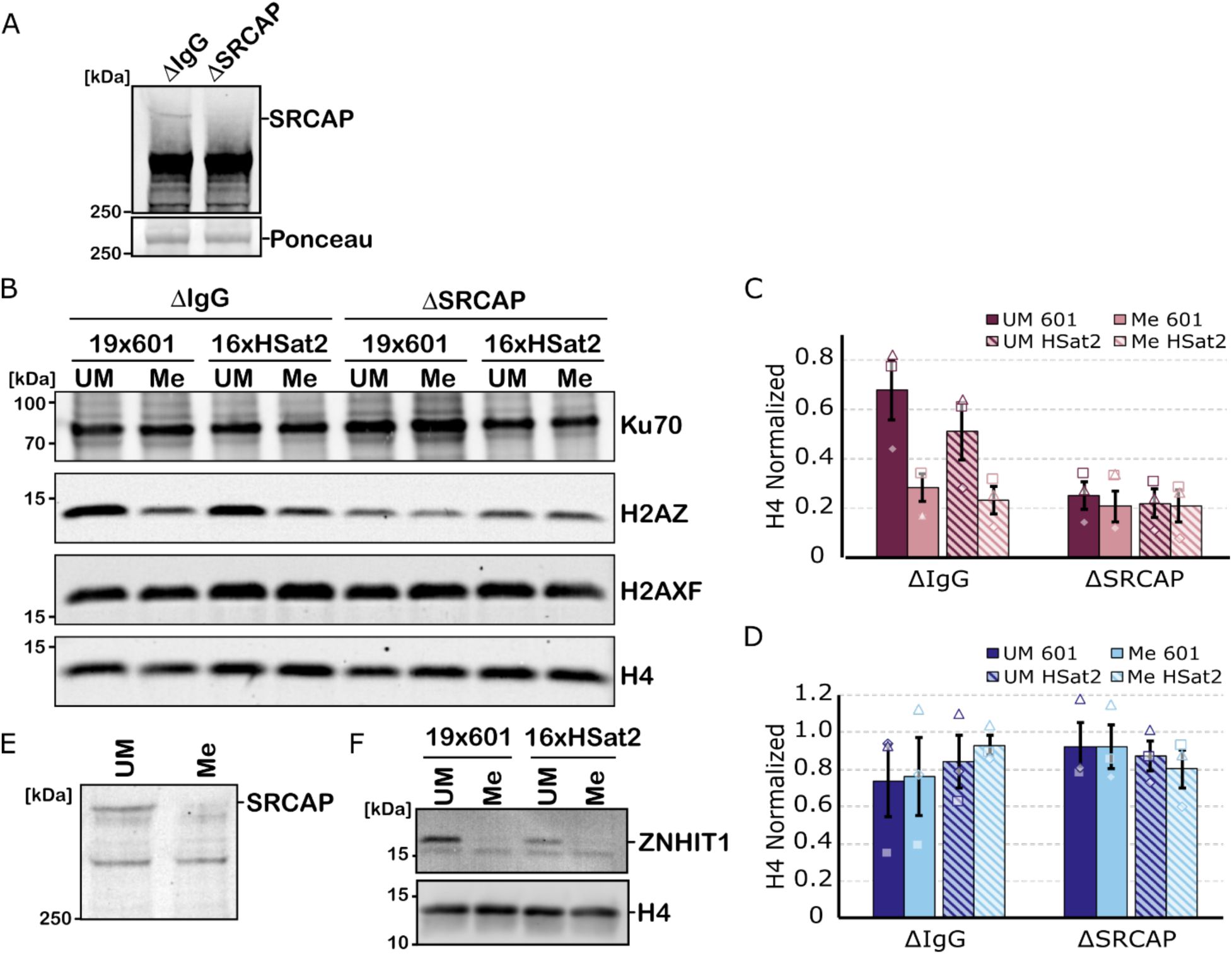
The SRCAP complex mediates H2A.Z’s preferential association with unmethylated DNA. **(A)** Western blot to detect SRCAP in control IgG-depleted *Xenopus* egg extract (ΔIgG) or SRCAP-depleted extract (ΔSRCAP). Bottom panel shows ponceau staining as a loading control. **(B)** Western blots of chromatinization assay in IgG control vs SRCAP depleted egg extract. Magnetic beads coated with 19-mer arrays of Widom 601 (19×601) or 16-mer arrays of HSat2 (16xHSat2) were incubated with interphase egg extract for 60 min, before their isolation. **(C)** Quantification of H2A.Z signal normalized to H4 from **(B)**. **(D)** Quantification of H2A.X-F signal normalized to H4 from **(B)**. Error bars in (**C**) and (**D**) represent SEM from n = 3 biological replicates and each shape represents data from one independent experiment. **(E)** Western blot staining for SRCAP on 16xHSat2 DNA beads incubated in interphase egg extract. **(F)** Western blot staining for ZNHIT1 on specified DNA beads incubated in interphase egg extract. Representative image shown from two independent experiments.

To understand the molecular basis of the SRCAP-C-dependent preferential deposition of H2A.Z to unmethylated DNA, we tested if DNA methylation inhibits SRCAP-C binding to DNA in *Xenopus* egg extracts. Indeed, SRCAP and ZNHIT1, two core subunits of SRCAP-C, were enriched on unmethylated but not on methylated DNA (**Fig 5E-F, Suppl Fig 13D**). In contrast, p400, the ATPase of TIP60-C, did not exhibit methylation-sensitive DNA binding (**Suppl Fig 15**). Since H2A.Z preferentially accumulates at TSSs, which tend to have more open/accessible chromatin, we explored the possibility that the apparent inhibition of SRCAP-C accumulation to methylated DNA was due to the potential capacity of DNA methylation to compact chromatin and change general DNA accessibility. If this were the case, we assumed that suppression of SRCAP-C binding to methylated DNA would not be detected if nucleosome assembly on DNA were to be inhibited. We tested this by depleting the HIRA complex, the major histone H3.3-H4 chaperone that is required for replication-uncoupled nucleosome assembly onto naked DNA in *Xenopus* egg extracts (Ray-Gallet et al., 2002), using antibodies against the HIRA complex subunit UBN2. Inconsistent with this hypothesis, SRCAP and ZNHIT1 still selectively bound to unmethylated DNA in UBN2-depleted extracts, where loading of H3, but not other DNA-binding proteins such as xKid and Dppa2 (Funabiki and Murray, 2000; Xue et al., 2013; Zierhut et al., 2014), was suppressed (**Suppl Fig 16**). These results indicate that exclusion of the H2A.Z chaperone SRCAP-C from methylated DNA is the major driver biasing H2A.Z deposition against methylated DNA.

## DISCUSSION

In this study, we described two distinct mechanisms via which DNA methylation can affect H2A.Z deposition and wrapping stability (**Fig 6**). We found that DNA methylation directly opposes H2A.Z deposition through inhibition of SRCAP-C DNA binding. SRCAP-C accounts for the majority of H2A.Z deposition on unmethylated DNA and is the primary mediator behind DNA methylation’s exclusion of H2A.Z in the transcriptionally silent *Xenopus* egg extract. A subpopulation of H2A.Z, however, is still deposited onto methylated DNA, driven by methylation insensitive mechanism(s) whose identity remains to be established. Once formed, H2A.Z nucleosomes on methylated DNA are more open and accessible compared to their unmethylated counterparts albeit the effect by DNA methylation is relatively subtle compared to that induced by H2A.Z incorporation. The impact on intrinsic nucleosome accessibility may also be dependent on wrapping/positioning ability of the DNA sequences used. This minor decrease in DNA-histone contact stability of methylated H2A.Z nucleosomes could conceivably contribute to clearance of H2A.Z from methylated regions of the genome, though such functional mechanisms remain to be established.

**Figure 6.**
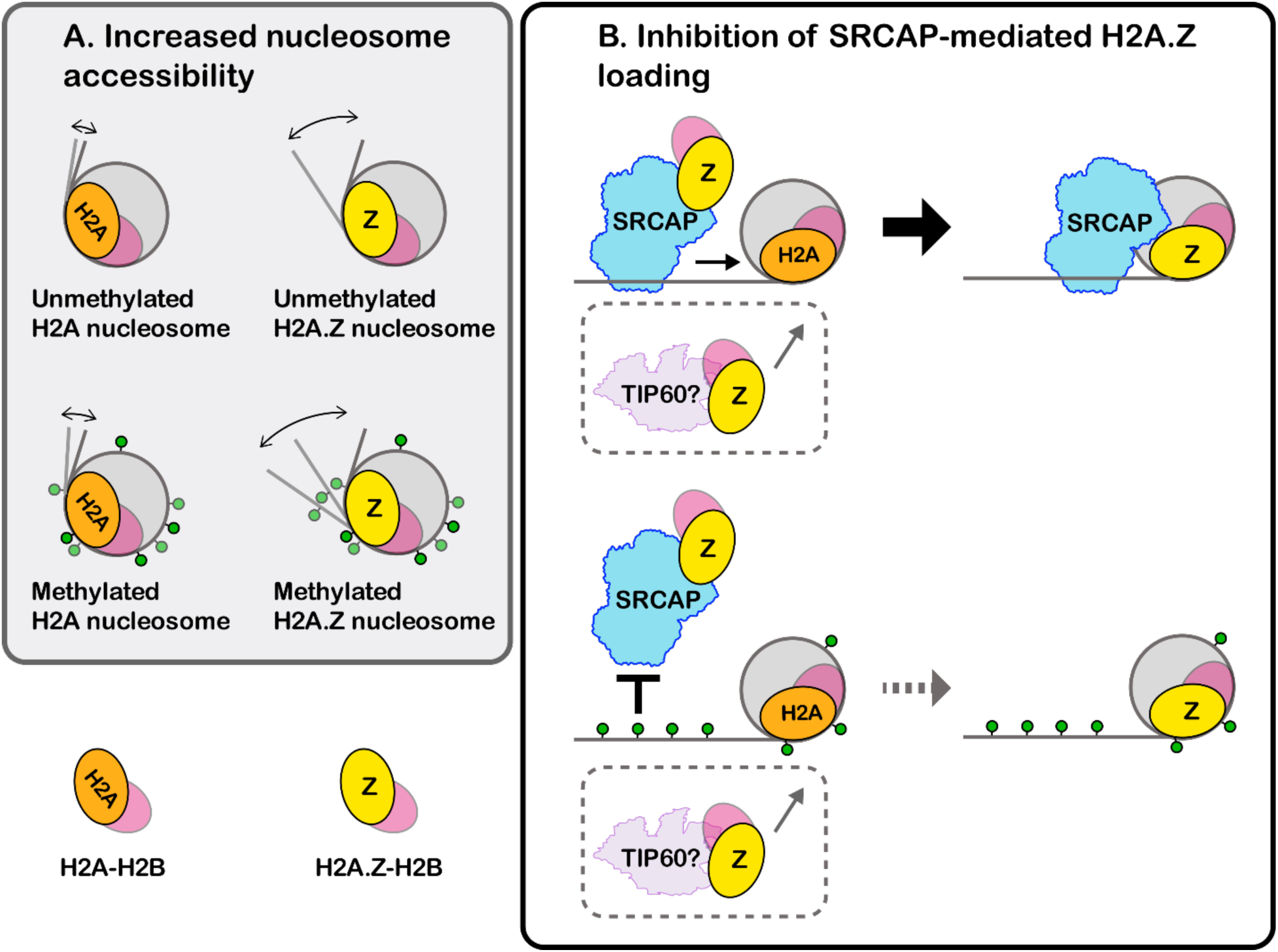
Schematic models for how DNA methylation influences physical nucleosome structure and SRCAP-mediated H2A.Z loading. (A) DNA methylation has little effect on nucleosome openness/accessibility of H2A nucleosomes. H2A.Z nucleosomes are more open/accessible than H2A nucleosomes, and DNA methylation DNA slightly opens H2A.Z nucleosome further causing nucleosomal DNA ends to be more accessible. (B) SRCAP-C is capable of binding to unmethylated DNA and replacing H2A with H2A.Z on the nucleosome. SRCAP-C cannot bind to methylated DNA, and thus H2A.Z deposition is suppressed on methylated DNA. An SRCAP-independent mechanism (possibly via TIP60) deposits H2A.Z in a DNA methylation insensitive manner.

Although a consensus on how DNA methylation influences nucleosome stability has yet to be achieved, our results from our cryo-EM analysis and endonuclease accessibility assay indicate that DNA methylation opens and increases accessibility of DNA segments particularly near the entry/exit region of the H2A.Z nucleosome core particle. This result is in line with reports based on FRET-based and molecular simulation studies on canonical H2A nucleosomes (Jimenez-Useche et al., 2014; Jimenez-Useche and Yuan, 2012; Li et al., 2022a; Ngo et al., 2016). Our *in silico* mixing 3D classification analysis indicates that more particles were classified as open and slid nucleosomes in the methylated H2A.Z sample than in unmethylated (**Figure 2E and F**), supporting the idea that DNA methylation destabilizes DNA-histone contacts. Since unmethylated G/C dinucleotides and methylated CpG are enriched at outward facing minor grooves and inward facing major grooves in the nucleosome core particles, respectively (Chodavarapu et al., 2010; Segal et al., 2006), DNA methylation may locally impact this DNA-histone contact stability. However, the ambiguity in DNA positioning of our cryo-EM structures prevents us from discussing the structural coordination of specific CpG sites. Alternatively, the increase in accessibility could be due to additive effects of DNA methylation on overall DNA conformation rather than at specific methylated segments. This is in agreement with observations that methylation increases DNA stiffness, making it harder for the DNA to bend and wrap around histone octamers (Ngo et al., 2016; Pérez et al., 2012).

Consistent with previous reports using the Widom 601 sequence (Jung et al., 2024; Lewis et al., 2021), our restriction enzyme accessibility assay shows that H2A.Z nucleosomes with HSat2 are considerably more open and accessible compared to canonical H2A, regardless of methylation status. This difference is substantially larger than that induced by DNA methylation on H2A.Z, suggesting that the physical influence of DNA methylation on nucleosomes is overall subtle. This apparent subtlety, however, may be due to the internal placement of the cut site that we tested, and thus the effect of DNA methylation was preferentially observed on H2A.Z nucleosomes, which are already highly accessible. Similar findings have been reported on canonical H2A 601 nucleosomes, where the effects of DNA methylation manifest greatest on an open nucleosome sub-population compared to their closed counterparts (Jimenez-Useche and Yuan, 2012). This is likely why we saw no methylation-dependent structural differences with the 601L sequence (**Suppl Fig 5**), as its innate nucleosome wrapping and positioning ability is even stronger than the classic Widom 601.

It is possible that the combined impact of DNA methylation and H2A.Z on nucleosome structure is further modulated by additional factors, including other histone variants, histone modifications, DNA sequences, and the position of DNA methylation. As H2A.Z frequently coexists with H3.3 at TSSs (Jin et al., 2009; Jin and Felsenfeld, 2007), H3.3 may conceivably contribute to H2A.Z’s sensitivity to DNA methylation, though it has been reported that H3.3 does not affect intrinsic H2A.Z nucleosome accessibility *in vitro* (Horikoshi et al., 2016; Jung et al., 2024; Thakar et al., 2009). Notably, in *Xenopus* extracts, nucleosomes assembled on naked DNA are exclusively composed of the H3.3 variant (Ray-Gallet et al., 2002), but the effect of DNA methylation on H2A.Z localization almost completely disappeared with SRCAP-C depletion (**Figure 5**). This indicates that the combination of H3.3 and H2A.Z alone is insufficient to explain suppression of H2A.Z loading to methylated DNA. Overall, we suggest that the subtle nucleosome destabilization effect of DNA methylation is preferentially pronounced on H2A.Z nucleosomes, which already exhibit an intrinsic wrapping instability compared to canonical H2A nucleosomes (**Figure 6**). Whether this subtle effect has functional significance remains a subject of future studies.

In *Xenopus* egg extract, H2A.Z deposition to exogenously added methylated DNA (both Widom 601 and HSat2) was about half that of their unmethylated counterparts (**Figure 5**), suggesting that the influence of DNA methylation on chaperone-mediated H2A.Z loading in egg extract is less sensitive to DNA sequence than its influence on nucleosome physical accessibility. Depleting SRCAP-C from *Xenopus* egg extracts effectively reduced, but not eliminated, H2A.Z deposition to nonmethylated DNA, but had little effect on the subpopulation of H2A.Z associated with methylated DNA. Therefore, *Xenopus* egg extracts appear to possess an additional SRCAP-C independent and DNA methylation-insensitive H2A.Z deposition mechanism (**Figure 6**). The known H2A.Z chaperone TIP60-C, whose DNA binding was not affected by methylation (**Suppl Fig 15**), is a prime candidate for this latter mechanism. Of note, concentrations of SRCAP and p400 (EP400), the ATPase subunits of SRCAP-C and TIP60-C, respectively, are comparable (13 nM v.s. 6 nM) (Wühr et al., 2014).

Exactly how SRCAP-C senses DNA methylation remains a pertinent question. Recent structural studies on the yeast SWR1 complex and human SRCAP-C indicate that both the ATPase and the ZNHIT1 subunit (Swc6 in yeast) mediate binding of, and one-dimensional diffusion on, naked DNA substrates (Louder et al., 2024; Park et al., 2024), marking them as likely candidates responsible for methylation sensing. Additionally, the YEATS domain-containing subunit GAS41, found in both SRCAP-C and TIP60-C, recognizes the open chromatin marks H3K14ac and H3K27ac (Hsu et al., 2018a, 2018b), and deposition of H3K27ac, itself, is antagonized by DNA methylation at enhancers (King et al., 2016). It is possible that both direct recognition of DNA methylation and changes in the local landscape of methylated chromatin play a role in dictating SRCAP-C localization and activity. However, SRCAP-C’s preference for unmethylated DNA was maintained even under nucleosome deficient conditions (**Suppl Fig 16**), indicating that SRCAP-C can sense DNA methylation status independent of chromatin accessibility and histone modifications.

Supporting our observation that a SRCAP-independent mechanism(s) exists to deposit H2A.Z on methylated DNA in *Xenopus* egg extracts, a substantial fraction (∼40 %) of H2A.Z-associated genomic DNA is methylated in sperm pronuclei. In the fibroblast cell line XTC-2, that methylated H2A.Z fraction is greatly reduced to 3 %. However, due to the accessibility bias inherent within CUT&Tag, our data likely underestimates the true degree of H2A.Z and DNA methylation overlap, particularly in the XTC-2 samples. For comparison, a previous study in human myometrium reported that ∼20 % of CpGs overlapping with H2A.Z are methylated, indicating that a sizeable population of colocalized H2A.Z and DNA methylation exists in somatic cells, which warrants further investigation (Berta et al., 2021). Nevertheless, we believe the decreased trend in H2A.Z and DNA methylation overlap of XTC-2 cells holds true and suggests that the high degree of H2A.Z and DNA methylation coincidence in sperm pronuclei reflects its unique early development and/or transcriptionally silent state. Given the strong preference of SRCAP-C binding to unmethylated DNA in *Xenopus* egg extracts, we predict that additional pathways missing in the egg extract system are necessary to reinforce the antagonism between DNA methylation and H2A.Z enrichment.

Perhaps the greatest difference between XTC-2 cells and *Xenopus* eggs is the presence of transcription, which is one of the main evictors of H2A.Z and necessary for global H2A.Z turnover as well as pruning of randomly deposited H2A.Z within gene bodies, where methylated cytosine enriches (Hardy et al., 2009; Lashgari et al., 2017; Ranjan et al., 2020; Tramantano et al., 2016). Traversing of RNA polymerase (Pol) II across nucleosomes, aided by the FACT complex, results in transient histone dimer displacement (Belotserkovskaya et al., 2003; Bevington and Boyes, 2013; Bintu et al., 2011; Kireeva et al., 2002), leading to the possibility that the more accessible H2A.Z nucleosome structures formed on methylated DNA may have increased susceptibility to Pol II disruption. Furthermore, FACT and Spt6 do not reincorporate H2A.Z into the reassembled nucleosome post Pol II encounter, which would reinforce selective loss of H2A.Z (Heo et al., 2008; Jeronimo et al., 2015). Outside of transcription, the egg extract system also lacks both the DNMT3 enzymes responsible for *de novo* methylation and the TET system responsible for DNA methylation removal in animals (Wühr et al., 2014). In *Arabidopsis*, H2A.Z promotes ROS1-associated DNA demethylation (Nie et al., 2019) and must be excluded to ensure faithful heterochromatin formation (Zhou et al., 2023). Thus, H2A.Z-driven mechanisms to limit DNA methylation remain relevant pathways to increase the H2A.Z/DNA methylation separation within the genome which may be missing in eggs/early embryonic states.

Beyond its interaction with DNA methylation, we show that H2A.Z promoter localization occurs immediately upon conditions mimicking fertilization in a transcription-independent manner. Unlike the almost complete loss of histones seen in mammalian sperm (Ward and Coffey, 1991; Wykes and Krawetz, 2003), *Xenopus* sperm is packaged as a mixture of protamines and H3/H4 tetramers with very low levels of H2A/H2B dimers, which are then remodeled into full nucleosomes upon exposure to the egg cytoplasm (Katagiri and Ohsumi, 1994; Mann et al., 1982; Oikawa et al., 2020; Shechter et al., 2009b). In our egg extract system, which lacks the maternal genome, we expect the bulk of H2A.Z in sperm pronuclei to be newly deposited and find that H2A.Z is still specifically targeted to promotors either via underlying sequence motifs (Ibarra-Morales et al., 2021; Murphy et al., 2018) or tetramer modifications maintained in sperm (Oikawa et al., 2020). This is similar to *Drosophila* where H2A.Z is recruited to TSSs prior to both Pol II binding and ZGA, marking genes for zygotic activation (Ibarra-Morales et al., 2021), indicating conservation of this initial transcription-independent H2A.Z positioning across both vertebrates and invertebrates.

Our work here provides a critical step in the pathways dictating genomic organization by uncovering the previously undescribed methylation sensitivity of SRCAP-C along with the subtle destabilizing effects of DNA methylation on H2A.Z nucleosomes (**Figure 6**). Together, these findings provide molecular mechanisms driving the conserved antagonism between H2A.Z and DNA methylation in the genome. However, many questions remain to be answered, such as the existence of both methylation sensitive and insensitive H2A.Z deposition pathways, the broader functional implications of the H2A.Z-DNA methylation antagonism as a whole, and mechanisms driving its maintenance in species that experience periods of global demethylation. These questions represent important areas of future study for our understanding of epigenome dynamics throughout development.

## MATERIALS AND METHODS

### Histone purification and refolding

pET3 vectors containing sequences for human histones H2A, H2A.Z, H2B, H3.2, and H4 (gifts from Shixin Liu) were expressed in Rosetta (DE3) cells (Novagene, Cat# 70954). Cells were pelleted 4000 x *g*, 20 min, 4 °C and resuspended in wash/lysis buffer [50 mM Tris-HCL pH 7.5, 100 mM NaCl, 1 mM EDTA, 5 mM β-mercaptoethanol, 0.2 mM PMSF] + 1 % Triton-X-100 then sonicated. Inclusion pellets were spun 25,000 x *g*, 20 min, 4 °C, and incubated with DMSO at room temperature (RT) for 30 min. Pellets were then resuspended in unfolding buffer [20 mM Tris-HCl pH 7.5, 7 M guanidine-HCl, 10 mM DTT], sonicated until pellets were fully broken up, and re-pelleted at 25,000 x *g*, 20 min, RT. The supernatant was then collected and dialyzed in urea dialysis buffer overnight at 4 °C [10 mM Tris-HCl pH 7.5, 7 M Urea, 1 mM EDTA, 5 mM β-mercaptoethanol, 100 mM or 200 mM NaCl for H2A/H2B and H3/H4, respectively]. Samples were then pelleted at 25,000 x *g*, 20 min, RT and the supernatant passed through Q Sepharose Fast Flow resin (Cytiva, Cat# 17051010). The flow-through was then collected and added to SP Sepharose Fast Flow resin (Cytiva, Cat# 17072910), washed with urea dialysis buffer, and eluted with elution buffer [10 mM Tris-HCl pH 7.5, 7 M Urea, 1 mM EDTA, 600 mM NaCl, 5 mM β-mercaptoethanol].

Equimolar ratios of histones for either octamer, or dimer and tetramer preparation were then mixed and dialyzed into refolding buffer [10 mM Tris-HCl pH 7.5, 1 mM EDTA, 5 mM β-mercaptoethanol, 2 M or 500 mM NaCl for octamers or dimers/tetramers, respectively]. Refolded histones were then purified via FPLC (ÄKTA pure, Cytiva, Cat# SN2626712) (Superdex 200 10/300 GL increase column, Cytiva, Cat# 28990944).

### Generation of DNA arrays

Generation of HSat2 arrays for use in chromatinization assays or arrays of Sat2R-P halves to prepare palindromic sequences for cryo-EM analysis was done as previously described in (Dyer et al., 2003). Briefly, DNA sequences of interest (see **Table 1**) containing one BglII site, one BamHI site, and two EcoRV sites surrounding the sequence were generated and purchased from Integrated DNA Technologies (IDT™) and cloned into the pUC18 vector just downstream of the XbaI site. The plasmid was then digested with either XbaI (NEB Cat# R0145) and BglII (NEB Cat# R0144) (to obtain the vector) or BglII and BamHI (NEB Cat# R3136) (to obtain the insert), and agarose gel purified. Vector and insert DNA were then ligated with T4 DNA ligase (NEB Cat# M0202). The digest and ligation steps were repeated until the desired repeat number was achieved.

**Table 1.**
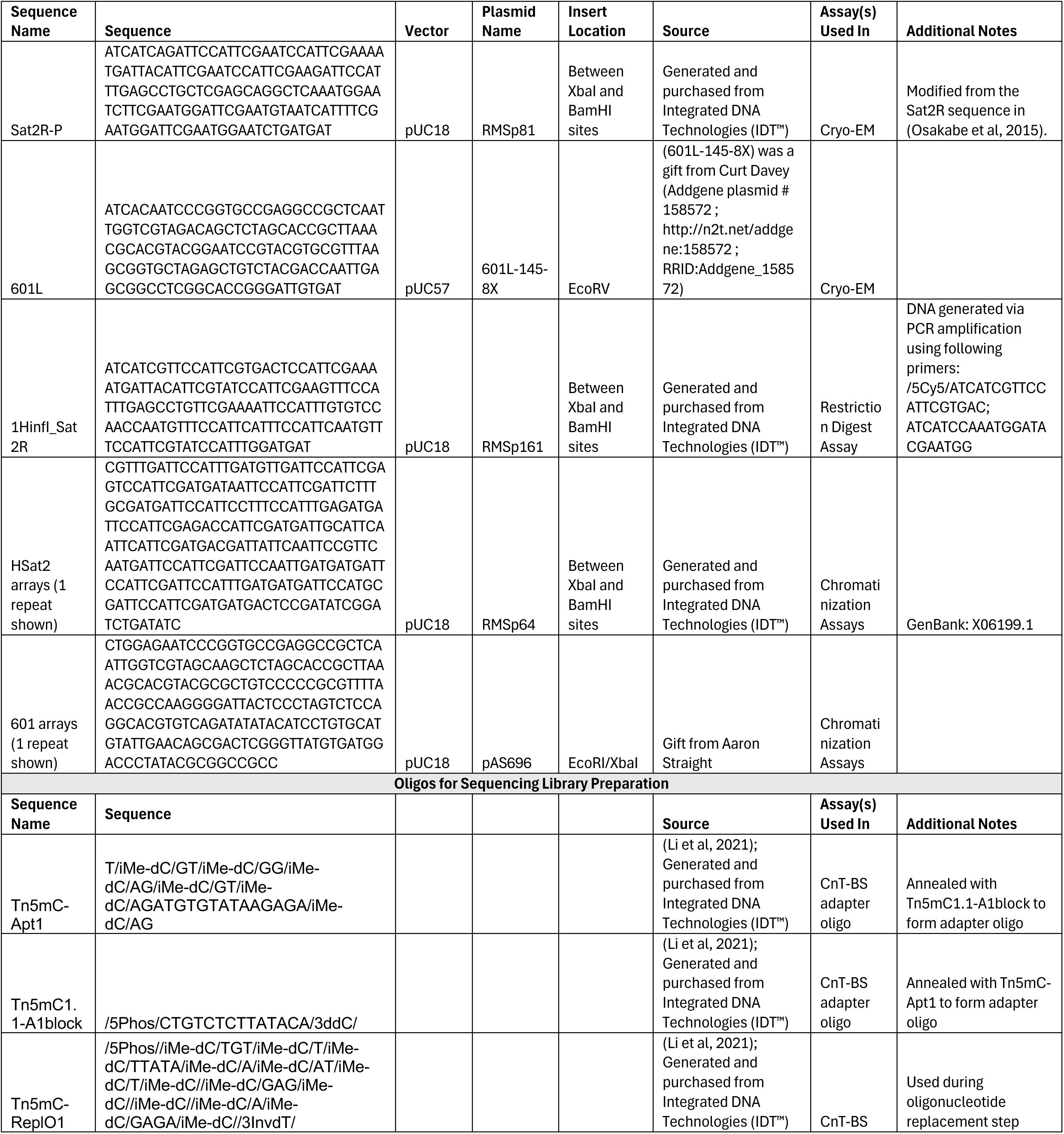
DNA sequences/oligos used in the study.

### Palindromic DNA preparation for Cryo-EM

Arrays of Sat2R-P halves were generated as described above. Plasmid containing an array of 601L halves (601L-145-8X) was a gift from Curt Davey (Addgene plasmid # 158572; http://n2t.net/addgene:158572; RRID: Addgene_158572) (Chua et al., 2012). Arrays of palindromic DNA halves for Cryo-EM analysis were then digested with EcoRV (NEB Cat# R3195) and purified via PEG precipitation (0.5 M NaCl, 8 % w/v PEG8000). Supernatant were collected and further purified via phenol-chloroform extraction. Insert ends were dephosphorylated with QuickCIP (NEB Cat# M0525) and phenol-chloroform extracted. Samples were then digested with either HinfI (NEB Cat# R0155) (601L DNA) or AvaI (NEB Cat# R0152) (Sat2R-P DNA), phenol-chloroform extracted, and FPLC purified in 1xTE buffer (Superdex 200 10/300 GL increase column, Cytiva, Cat# 28990944). DNA halves were then incubated with T4 DNA ligase (NEB Cat# M0202) per manufacturer’s protocol until all halves were completely ligated.

### Methylation of DNA substrates

DNA substrates were enzymatically methylated using the *M.SssI* CpG methyltransferase (NEB, Cat# M0226). Approximately 160 U of methyltransferase were used for 50 µg of DNA in 1x NEB 2 buffer with 500 µM SAM. Samples were incubated at 37 °C for at least one hour. Completion of methylation was checked using the methylation sensitive restriction enzyme BstUI (NEB, Cat# R0518). DNA was purified using phenol-chloroform extraction prior to use.

### Nucleosome Reconstitution

DNA and either histone octamers or dimers and tetramers were mixed at a final target concentration of 1 µM in 1x TE buffer and 2 M NaCl. Exact ratios of DNA to histones were titrated for each reconstitution batch, starting at equimolar ratios. Samples loaded in dialysis buttons were then added to high salt dialysis buffer [10 mM Tris-HCl pH 7.5 at 4 °C, 1 mM EDTA, 2 M NaCl, 0.01 % Triton X-100, 5 mM β-mercaptoethanol] at 4 °C. The high salt dialysis buffer was then exchanged into low salt dialysis buffer [10 mM Tris-HCl pH 7.5 at 4°C, 1 mM EDTA, 50 mM NaCl, 0.01 % Triton X-100, 5 mM β-mercaptoethanol] at a rate of ∼0.7 ml/min over a period of 30-40 h. Nucleosome quality was assessed on a 5 % 37.5:1 acrylamide/bis 0.5 x TBE gel (120 V).

### Mass Photometry Analysis of Nucleosomes

Mass photometry analysis was performed on a Refeyn Two^MP^ instrument and analyzed using DiscoverMP v2022R1 (with IScat v1.52.0 and IScat Utils v1.43.0). 24 x 50 mm with 170 ± 5 µm thickness No. 1.5H glass slides (ThorLabs, Cat# CG15KH1) and silicone gaskets (Grace Bio-labs, Cat# 103250) were washed 2x with alternating 100 % isopropanol and deionized water. Clean gaskets were then placed on the cleaned slides to serve as a sample chamber. 5 µl of 1x PBS was placed in the gasket to calibrate each sample collection then 5 µl of nucleosome samples were added for a target final concentration of ∼5-10 ng/µl nucleosomes in the well. Measurements were conducted at room temperature.

### Cryo-EM Sample Preparation and Data Collection

Human H2A.Z nucleosomes were reconstituted on either 601L or Sat2R-P DNA as described above with a final concentration aim of 1.6 µM in 1 ml. Reconstituted nucleosomes were dialyzed into buffer containing 10 mM HEPES-KOH pH 7.4 and 30 mM KCl then concentrated to a final concentration of ∼1 mg/ml using Amicon® Ultra Centrifugal Filters (Cat# UFC5010). Samples were then mixed with 10x cryo-protectant buffer [10 mM HEPES-KOH pH 7.5, 30 mM KCl, 10 % Trehalose, 1 % Hexanediol, 1 mM MgCl_2_] and 3 µl of each solution was applied to plasma-cleaned) Quantifoil gold R1.2/1.3 400-mesh grids (Quantifoil) either graphene coated (graphene coating was done in-house as specified in (Arimura et al., 2025) or not for the Sat2R-P and 601L samples, respectively. Grids were then vitrified using the Vitrobot Mark IV (FEI) at 100 % humidity, 2 sec blotting time, blot force 1. Data collection was conducted using the Titan Krios (ThermoFisher), equipped with a 300 kV field emission gun and a K3 direct electron detector (Gatan), at a magnification of x81,000 (0.86 Å/pixel). Total micrographs collected for each sample as follows: 4544 for unmethylated Sat2R-P nucleosomes; 6891 for methylated Sat2R-P; 7020 for unmethylated 601L; 5427 for methylated 601L. Data statistics summarized in **Suppl Table 1**.

### Cryo-EM Data Analysis

Diagrams of analysis workflows are shown in **Suppl Fig 1** and **Suppl Fig 5**. Micrographs were motion-corrected using MotionCor2 (Zheng et al., 2017) in Relion v4 (Scheres, 2012). Downstream analysis was conducted in CryoSPARC v4.6 (Structura Biotechnology Inc.). Motion-corrected micrographs then underwent patch CTF correction. Particles were picked using Topaz v0.2 (Bepler et al., 2019) trained on a set of 1,000-2,000 manually picked nucleosome-like particles and then extracted to 384 pix (∼330 Å) boxes with 4x binning and used for 2D classification into 200-250 classes. Particles from selected classes were then run through a series of heterogeneous refinements with 1-2 nucleosome-like models and 4 decoy models generated through *ab initio* reconstruction to sort out junk particles from those containing nucleosomes. The heterogenous refinements were iterated until greater than 90 % of particles in the round were classified as nucleosome-like. These particles were then further refined either via Refine 3D, CTF refinement, and polishing in Relion v4 (601L structures) or homogeneous and non-uniform refinements in CryoSPARC v4.6 (Sat2R-P). Particles for the methylated Sat2R-P structure underwent another heterogeneous refinement using a closed nucleosome, open, and hexasome model as starting classes to select for the highest resolved closed nucleosome structure. Final resolutions were determined using the gold standard FSC threshold (FSC = 0.143). Local resolution maps shown in **Suppl Fig 3** were generated using the local resolution estimation tool with default parameters in CryoSPARC. The maps were then visualized on density maps filtered using the local filter tool with default parameters. For the *in silico* mixing 3D classification analysis (**Suppl Fig 4**): micrographs from both unmethylated and methylated Sat2R-P structures were merged and subjected to the data analysis pipeline described above. Six starting models representing different nucleosome structure types were then constructed using the ab initio 3D reconstruction tool. These models were then used as input volumes, and cleaned particles from the merged analysis were then sorted to each of these models using the heterorefinement tool in CryoSPARC.

### Atomic Modeling and RMSD analysis

Initial atomic models were generated using previously published structures of human H2A.Z-containing histone core (PDB ID: 3WA9) (Horikoshi et al., 2013) and modified versions of either 601L DNA (PDB ID: 3UT9) (Chua et al., 2012) or Sat2R DNA (PDB ID: 5CPI) (Osakabe et al., 2015). Models were docked into the EM density maps using Phenix v1.21, then iteratively refined and assessed in Phenix v1.21 (Liebschner et al., 2019) and *Coot* 0.9.8.7 (Emsley et al., 2010). RMSD values were calculated in PyMOL v2.5.7 (Schrödinger) using the cealign function. For the Sat2R-P models, DNA positioning was determined using the EM densities at the dyad of the unmethylated map which allowed for pyrimidine and purine distinction. A two-base pair shift was determined by assessing ambiguity of densities around nucleotides which change from purine to pyrimidine, and *vice versa*, between mirrored DNA models versus those which retain identity. After determining best-fit DNA models for the unmethylated structure, the same DNA positionings were then used for the methylated structure.

### DNA Preparation for Restriction Digest Assay

Non-palindromic 1HinfI_Sat2R DNA (**Table 1**) was prepared via PCR using Q5 polymerase (NEB Cat# M0491). Final concentrations of reaction mixture as follows: 1x Q5 reaction buffer; 200 µM dNTPs; 2 µM forward and reverse primers; 0.2 g/µl DNA template; 0.02 U/µl Q5 polymerase. Roughly 60-65 PCR cycles were run following manufacturer’s protocol with a melting temperature of 60 °C. PCR products were phenol-chloroform extracted and then FPLC purified in 1xTE buffer (Superdex 200 10/300 GL increase column, Cytiva, Cat# 28990944).

### Restriction Digest Assay

Nucleosomes were mixed at 30 ng per timepoint sample in 1x rCutSmart® buffer (NEB Cat# B6004S) [50 mM potassium acetate, 20 mM Tris-acetate pH 7.9 at RT, 10 mM magnesium acetate, 100 µg/ml recombinant albumin] and 0.5 U/µl HinfI restriction enzyme (NEB Cat# R0155S). Samples were incubated at 37 °C and added to stop buffer [final concentrations: 20 mM Tris-HCl pH 8.0 at RT, 20 mM EDTA, 0.1 % SDS, 0.6 mg/ml Proteinase K] at each timepoint. Stopped samples were then incubated at 55 °C for 40 min and run on a 10 % 37.5:1 acrylamide/bis 0.5x TBE gel (55 min at 150 V) and stained with SYBR™ Safe DNA stain (Thermo Fisher Scientific, Cat# S33102). Fluorescent signals were measured on a LI-COR Odyssey M system and quantified using ImageJ via histogram analysis as previously described in (Stael et al., 2022). The ratio of Cy5: SYBR™ Safe signals was then determined for each time point and plotted. A two-tailed t-test assuming unequal variance was then performed on data from n = 5 experiments for statistical analysis.

### Antibodies

Antibodies used in this study are listed in **Table 2**. Custom antibodies against *Xenopus laevis* H2A.Z, SRCAP, and UBN2 were generated through Cocalico Biologicals, Inc.™ using their standard protocol. Target binding of the H2A.Z antibody used for CnT-BS and SRCAP custom antibody in *Xenopus* egg extract was verified using mass spectrometry analysis (MS) (**Supplementary Data 1 and 2**).

**Table 2.** Antibodies used in this study.

| Antigen | Host | Company | Catalog No. | Peptide Used for Generation | Dilution | Assay(s) Used In |
| --- | --- | --- | --- | --- | --- | --- |
| XI H2A.Z | Rabbit | Custom |  | CIHKSLIGKKGQQKTV | 2 µg/ml | Western Blots |
| H2A.Z | Rabbit | Active Motif | Cat#39013 |  | 1:10 | CnT-BS |
| H3 | Mouse | Cosmo Bio USA | MABI0001-100-EX |  | 1:100 | CnT-BS |
| H4 | Mouse | Active Motif | Cat# 61521 |  | 1:1000 | Western Blots |
| XI H2A.X-F | Rabbit | Custom, gift from David Shechter |  | Shechter et al. 2009 | 1:1000 (of serum) | Western Blots |
| XI Ku70 | Rabbit | Custom |  | Postow et al. 2008 | 2 µg/ml | Western Blots |
| XI SRCAP | Rabbit | Custom |  | CPNSPSSHRVRKAKT | 0.8 µg per 1 µl of beads (depletions);<br>12 µg/ml (WB) | Depletions;<br>Western Blots |
| hsSRCAP | Rabbit | Kerafast | ESL103 |  | 0.8 µg per 1 µl of beads (depletions);<br>1:500 (WB) | Depletions;<br>Western Blots |
| ZNHIT1 | Rabbit | Proteintech | 16595-1-AP |  | 1:1000 | Western Blots |
| P400 | Rabbit | Bethyl | A300-541A |  | 1:1000 | Western Blots |
| ALFA tag | Alpaca (single domain antibody purified from E. coli) | NanoTag Biotechnologies | N1502-Li800-L |  | 1:1000 | Western Blots |
| XI UBN2 | Rabbit | Custom |  | CGGQNKGDAKLPRKNR | 1.5 µg per 1 µl of beads (depletions);<br>2 µg/mL (WB) | Depletions;<br>Western Blots |
| xKID | Rabbit | Custom |  | Funabiki and Murray 2000 | 1 µg/mL | Western Blots |
| XI Dppa2 | Rabbit | Custom |  | Xue et al. 2013 | 5 µg/mL | Western Blots |
| **Secondaries** |  |  |  |  |  |  |
| IRDye® 800CW anti-Rabbit IgG | Goat | LICORbio | 926-32211 |  | 1:10000 | Western Blots |
| IRDye® 680RD anti-Mouse IgG | Goat | LICORbio | 926-68070 |  | 1:10000 | Western Blots |
| anti-Rabbit IgG | Guinea Pig | antibodies-online.com | ABIN101961 |  | 1:100 | CnT-BS |
| anti-Mouse IgG | Rabbit | abcam | ab46540 |  | 1:100 | CnT-BS |

### Mass Spectrometry Validation

For the MS analysis, 20 µl of antibody-bound Protein A Dynabeads™ (Invitrogen, Cat# 10008D) (0.8 µg antibody per 1 µl of beads) were incubated at a 1:1 ratio with CSF egg extract (see next section) at 4 °C for 1h. Samples were collected via magnet and washed 3 times with sperm dilution buffer (SDB) [150 mM Sucrose, 10 mM HEPES-KOH pH 8.0, 1 mM MgCl_2_, 100 mM KCl] then run on a 4-20% gradient SDS-PAGE gel as a 1 cm short-stack. The gel was then stained with Coomassie Brilliant Blue G-250 (Thermo Fisher) and the short stack was cut out. The cut gel was then destained using a solution of 30 % acetonitrile and 100 mM ammonium bicarbonate in water and dehydrated using 100 % acetonitrile. Samples were subjected to heat assisted (50 °C) trypsinization for 1 h. The resulting peptides were extracted three times with a solution of 70 % acetonitrile and 0.1 % formic acid then concentrated using a SpeedVac and resuspended in 10 µl volumes. 3 µl of purified peptides were subsequently analyzed by LC-MS/MS using a Dionex 3000 HPLC system equipped with an EasySPrayer (ES902), coupled to an Orbitrap ASCEND mass spectrometer from Thermo Scientific. Peptides were separated by reversed-phase chromatography using solvent A (0.1 % formic acid in water) and solvent B (80 % acetonitrile, 0.1 % formic acid in water) across a 45-min gradient. Data was processed using ProteomeDiscoverer 1.4 and quantified using ProteomeDiscoverer v1.4/Mascot v3.1. The spectra were then queried against a *Xenopus laevis* database (Wühr et al., 2014) and matched peptides filtered using a Percolator calculated FDR of 1 %. As proxy for protein amount, the average area of the three most abundant peptide signals was used. Relative enrichment of proteins in the sample was then calculated by dividing by known protein concentrations in *Xenopus laevis* eggs and then the enrichment over IgG control was calculated. Common contaminates were marked in red.

### *Xenopus laevis* frogs and preparation of sperm and CSF egg extract

Female *Xenopus laevis* frogs were purchased from Xenopus 1. Male frogs were purchased from either Xenopus 1 (for use in non-sequencing experiments) or the National Xenopus Resource (*XLa.J-Strain^NXR^,* Cat# NXR_0024; for use in sequencing experiments). Vertebrate animal protocols (20031 and 23020) approved by the Rockefeller University Institutional Animal Care and Use Committee were followed for housing and care.

CSF egg extract and demembranated sperm nuclei were prepared as specified previously (Tutter and Walter, 2006).

### Sperm Pronuclei Preparation

Sperm from J-strain frogs were added to prepared CSF egg extract at a final concentration of 1,500 sperm/µl extract with a total extract volume of 200 µl for each sample. 90 ng (∼1 % of total DNA) of lambda DNA (Promega Cat# D1521) was also added to assess bisulfite conversion efficiency and the extract solution was then cycled to interphase with the addition of CaCl_2_ to a final concentration of 0.3 mM and incubated at 20 °C for 90 min. Samples were then diluted 1:4 with 1xDB3 buffer [10 mM HEPES-KOH pH 8.0, 100mM NaCl, 0.5 mM spermine, 0.5 mM spermidine, 1 mM ASEBF (GoldBio Cat#A-540-1), 1x ROCHE cOmplete™ protease inhibitor cocktail (Cat# 11697498001), 200 mM Sucrose, 0.1 % Tween-20], layered onto 1 ml of 60SDB3 [1xDB3 + 60 % sucrose] and centrifuged at 4 °C, 10,000 rpm for 45 min (Beckman Coulter SX241.5 rotor). Sample pellets were washed 2x with 1xDB3, centrifuging at 4,000 x *g*, 4 °C for 4 min, then resuspended in 1x CnT-WB [20 mM HEPES-KOH pH 7.8, 150 mM NaCl, 0.5 mM spermidine, 1x ROCHE cOmplete™ protease inhibitor cocktail, 0.1 % Tween-20].

### XTC-2 Cell Line

XTC-2 cells (gift from Dr. David Shechter) were cultured in Leibovitz’s L-15 Medium (Gibco™, Cat# 11415064) (0.7x amphibian strength, diluted with cell culture grade water (Corning®, Cat# 46-000-CV)) supplemented with 500 U/ml each of penicillin and streptomycin (Gibco™, Cat# 15140122), and 10 % FBS (R&D systems, Cat# S11150). XTC-2 cells were cultured in a 26 °C incubator without CO_2_ supply. Cell line identity was verified via mitotic-spread karyotyping.

### XTC-2 Nuclei Preparation

XTC-2 cells were harvested at 90-100 % confluency from a 15 cm dish. Nuclei isolation was performed at 4 °C using the Nuclei EZ Prep kit (Millipore Cat# NUC101-1KT) as per manufacturer’s protocol and resuspended in 1x CnT-WB. For each sample, 18 ng (∼1 % of total DNA) of unmethylated lambda DNA (Promega Cat# D1521) was added to 300,000 nuclei to assess for bisulfite conversion efficiency.

### Purification of Tn5

Tn5 was purified as described in (Soroczynski et al., 2024). Briefly, VR124 pETv2-10His-pG-Tn5 (E54K, L372P) *E. coli* strain (Addgene #198468) were cultured to log growth phase and Tn5 expression induced with IPTG addition. Cells were harvested at 8,000 x *g*, 30 min at 4 °C, washed with ice-cold DPBS, re-pelleted and frozen. Frozen pellets were then resuspended at 4 °C in lysis buffer LysEQ [20 mM HEPES pH 7.8, 800 mM NaCl, 20 mM imidazole, 10 % glycerol, 1 mM EDTA, 2 mM TCEP-HCl, 1x EDTA-free cOmplete protease inhibitor] and sonicated. Lysates were pelleted at 20,000 x *g*, 35 min at 4 °C, then filtered through a 0.45 µm PES filter. Supernatant was then loaded onto a HisTrap HP 5 ml column (Cytiva, Cat# 17-5247-01) pre-equilibrated with LysEQ buffer. The loaded column was then washed with 10 column volumes each of WashB1 [20 mM HEPES pH 7.8, 800 mM NaCl, 30 mM imidazole, 10 % glycerol, 1 mM EDTA, 2 mM TCEP-HCl] and WashB2 [20 mM HEPES pH 7.8, 800 mM NaCl, 45 mM Imidazole, 10 % glycerol, 1 mM EDTA, 2 mM TCEP-HCl]. Protein was then eluted with EluB [20 mM HEPES pH 7.8, 800 mM NaCl, 400 mM imidazole, 10 % glycerol, 1 mM EDTA, 2 mM TCEP-HCl]. Tn5-containing fractions were pooled and dialyzed 4 °C overnight in SECB buffer [50 mM Tris-HCl pH 7.5, 800 mM NaCl, 10 % glycerol, 0.2 mM EDTA, 2 mM DTT] then filtered through a 0.45 µm PES filter and FPLC purified on a HiLoad 16/600 Superdex 200 pg column (Cytiva, Cat#: 28989335). Purified fractions were pooled and concentrated with a 30 MWCO Amicon Ultra Centrifugal filter (final concentration 12 µM) and mixed with UltraPure glycerol to a final concentration of 55 % glycerol. Purified Tn5 was stored at −20 °C.

### CUT&Tag-Bisulfite Library Preparation

Sequencing libraries were prepared as previously described in (Li et al., 2021). Briefly, prepared nuclei samples were bound to 10 µl of activated concanavalin A beads (EpiCypher Cat# 21-1401) and incubated with specified primary and secondary antibodies (**Table 2**) in 50 µl Antibody Binding Buffer [CnT-WB + 2 mM EDTA + 0.1 % w/v BSA] for 1 h shaking at RT, washing 3x with CnT-WB between each incubation. Samples were then incubated with 50 µl of transposome complex (1:1 pG-Tn5:adaptor primer mix, ∼0.1-0.2 µM final to each sample) in CnT-300 [CnT-WB with 300 mM NaCl] for 1 h at RT then washed with CnT-300. Transposome activity was initiated with addition of 300 µl Tagmentation Buffer [20 mM HEPES-KOH pH 7.8, 300 mM NaCl, 0.5 mM spermidine, 10 mM MgCl_2_, 1x ROCHE cOmplete™ protease inhibitor cocktail, 0.1 % Tween-20] and incubated 37 °C, 1500 rpm on thermomixer, 1 h. Reactions were stopped with addition of stop buffer [15 mM EDTA, 0.1 % SDS, 0.2 mg/mL Proteinase K final concentrations] and incubated at 55 °C overnight, then phenol-chloroform extracted. Purified samples were resuspended in 11 µl of 10 mM Tris-HCl, pH 8.0 and oligonucleotide replacement/gap repaired carried out through addition of 2 µl of 10 µM Tn5mC-ReplO1 DNA oligo (**Table 1**), 2 µl 10x Ampligase Buffer (Lucigen), 2 µl dNTP mix (2.5 mM for each dNTP). Solutions were subject to the following heating protocol: 50 °C 1 min, 45 °C 10 min, cool to 37 °C at −0.1 °C/second. Once at 37 °C, 3U of T4 DNA polymerase and 12.5U of Ampligase (Lucigen) were added and samples incubated for 30 min. Reactions were stopped with 20 mM EDTA final and purified using the MinElute PCR purification kit (Qiagen Cat# 28004). Samples were bisulfite converted using the EZ DNA Methylation-Lightning Kit (Zymo Research Cat# D5030) and libraries amplified (14 cycles) using NEBNext Q5U Master Mix (NEB Cat# M0597) and Illumina Nextra i5/i7 adapter primers.

### Sequencing Data Analysis

#### Adapter trimming, deduplication, and genome alignment

Libraries were sent to Novogene for sequencing on the NovaSeqX Plus system (PE150). XTC-2 data was then downsampled to 30 million reads using seqtk (v1.5-r133) to have all samples at similar read depths. Adapter trimming and dedeuplication of the demultiplexed reads was then done using fastp v0.24.0 (Chen, 2023) [--adapter_sequence CTGTCTCTTATACAC --adapter_sequence_r2 CTGTCTCTTATACAC -g --poly_g_min_len 5 -p -l 20 --cut_tail -y -Y 20 -q 20 –dedup]. Processed reads were then aligned to the *Xenopus laevis* genome (Xenla 10.1) and methylated CpG proportions for all mapped reads calculated using Bismark v0.24.2 (Krueger and Andrews, 2011) [-q --local --score-min G,20,8 --ignore-quals --no-mixed --no-discordant --dovetail --maxins 700].

#### Count matrix generation and sequencing depth normalization

The *Xenopus laevis* genome was subset into 1000 bp bins. Reads from processed bam files were then tallied according to bins to generate a genome-wide count matrix. Mitochondrial associated reads and bins which contained no reads across all samples were removed. Size factors were then estimated using DESeq2 v1.42.1 (Love et al., 2014) to generate sequence depth normalized bigwig files. Count matrices were also vst transformed in DESeq2 then used to generate the read fragment correlation plot. For other analyses, only genomic bins containing at least 17 reads across at least 2 samples were kept and replicates for each condition were averaged.

#### DNA Methylation analysis

To get methylated CpG proportions for each sample (**Fig 4A**), H2A.Z peaks were first identified with SEACR (v1.3) (Meers et al., 2019) using a cutoff of 0.025 by AUC with a stringent threshold. Individual CpG data was then extracted using Bismark with a cutoff of 5 reads and disregarding the first 9 bp of read 2 (due to adapter fill-in). Methylation proportions for H2A.Z samples were then calculated for CpGs overlapping with H2A.Z peaks. For H3, only regions outside of identified H2A.Z peaks were used to assess the CpG methylation status of general genomic background non-coincident with H2A.Z enriched areas. To generate H3-associated methylation 500 bp tiles for use in plotting (**Suppl Fig 9B-D**), methylation data was extracted with a specified cutoff of 1 read and disregarding the first 9 bp of read 2 then imported into a methylRawList object with a minimum read coverage of 2 and counted over 500 bp tiles with a 500 bp step size and a minimum of 2 CpG sites using methylKit v1.28.0 (Akalin et al., 2012).

#### Generation of plots

Genomic annotations for the top 20 % of filtered bins containing the most reads for each condition were generated and proportions plotted using ChIPseeker v1.38.0 (Yu et al., 2015) with reference gene annotations gathered from Xenbase (Fisher et al., 2023). Heat maps and plot profiles were created using deepTools v3.5.5 (Ramírez et al., 2016). For comparison of H3 methylation sites and H2A.Z, a signal matrix was first generated using the tiled H3 methylation bigwigs against all annotated *Xenopus laevis* genes, removing genes with no methylation data available. The resulting gene list was then used to plot corresponding H2A.Z signals.

### *in vitro* mRNA Synthesis

DNA sequences of proteins of interest were cloned into pSP64T plasmids modified to include a Kozak sequence (GCCACC) upstream of the coding sequence and linearized. pSP64T (DM#111) was a gift from Douglas Melton (Addgene Plasmid# 15030). RNA production was carried out using the MEGAscript™ SP6 transcription kit (Invitrogen, Cat# AM1330) as per manufacturer’s protocol, with the addition of 4.5 mM CleanCap® Reagent AG (TriLink, Cat# N-7113) and a reduction in GTP concentration down to 1.5 mM final.

### Chromatinization in Egg Extract Assays

pAS696 containing a 19-mer array of Widom 601 DNA was a gift from Aaron Straight (Guse et al., 2011). 601 and HSat2 arrays were digested with XbaI and EcoRI or XbaI and BamHI, respectively, along with DraI and HaeII. DNA fragments containing the array sequence were then purified through a PEG precipitation titration, starting at 4.5 % of PEG (MW8000) and increasing in 0.5 % increments until 8 % (NaCl added to 0.5 M final concentration in the starting solution). Fragment containing fractions were then assessed via agarose gel and purified via phenol-chloroform extraction. Fragment ends were biotinylated with biotin-14-dATP (Jena Bioscience, Cat# NU-835-BIO14-S) using Klenow Fragment (3’→ 5’ exo-) (NEB Cat# M0212) following manufacturer’s protocol. Biotinylated DNA was phenol-chloroform extracted and bound to M-280 Streptavidin Dynabeads™ (Invitrogen, Cat# 11206D) at 300 ng DNA per 1 µl of beads. DNA bead binding was carried out at room temperature for a minimum of 1 h shaking in the following solution: 1x DNA Bead Binding buffer (DBB) [750 mM NaCl, 50 mM Tris-HCl pH 8.0, 0.25 mM EDTA, 0.05 % Triton-X-100]; 2.5 % (w/v) polyvinyl alcohol; 750 mM NaCl. DNA beads were then washed 2x with DBB and 3x with sperm dilution buffer (SDB) [150 mM Sucrose, 10 mM HEPES-KOH pH 8.0, 1 mM MgCl_2_, 100 mM KCl] and added to egg extract at a ratio of 4 µl DNA beads to 30 µl of egg extract. Experiments that did not use mRNA for exogenous protein expression also included 0.1 mg/ml cycloheximide dissolved in EtOH. CaCl_2_ was then added to the extracts at a final concentration of 0.3 mM and incubated at 20 °C for 1 h. Samples were then diluted 10-fold with SDB, and DNA beads were collected on a magnetic rack and washed 3x with SDB + 0.05 % Triton-X-100 before running on SDS-PAGE gel and assessed via western blot.

### Western Blot Analysis

Samples were boiled in 1x SDS loading buffer [125 mM Tris-HCl pH 6.8, 0.01 % SDS, 25 % glycerol, 0.03 mg/ml bromo-phenol blue, 350 mM 2-mercaptoethanol] then run on either a 4-20 % SDS-PAGE gel (Bio-Rad, Cat# 4561096) or 3-8 % Tris-acetate gel (for SRCAP detection) (Invitrogen, Cat# EA0375BOX). Samples were then transferred from the gel to a nitrocellulose membrane (Cytiva, 0.1 µm pore size, Cat# 10600000) overnight at 4 °C in 1x transfer buffer [20 mM Tris, 150 mM Glycine, 20 % Methanol] (14V). Membranes were blocked for 30 min at RT with Intercept® (TBS) Protein-Free Blocking Buffer (LICORbio, Cat# 927-80001), followed by primary antibody incubation (diluted in Intercept® T20 (TBS) Antibody diluent (LICORbio, Cat# 927-65001)) for 1 h at RT. Membranes were then washed 3x, 5 min each wash, in 1xTBS+0.05 % Tween-20 then incubated with appropriate secondary antibodies for 1 h at RT and washed. Blots were then imaged on a LI-COR Odyssey M system and signals quantified using ImageJ via histogram analysis as previously described in (Stael et al., 2022). Ratios of either H2A.Z or H2A.X-F signals to H4 signals were then determined for normalization purposes in the chromatinization assays.

### Antibody Depletion from Egg Extract

Either SRCAP or UBN2 antibody was bound to Protein A Dynabeads™ (Invitrogen, Cat# 10008D) at a concentration of either 0.8 µg or 1.5 µg antibody per 1 µl of beads, respectively, overnight at 4 °C in wash/coupling (W/C) buffer [10 mM HEPES-KOH pH 8.0, 150 mM NaCl]. Beads were then washed 2x with W/C buffer and 3x with SDB. An equal volume of beads and egg extract (supplemented with 0.1 mg/ml cycloheximide) were then incubated together for 1 h at 4 °C and beads removed via magnet.

### Other Instruments and Software Used

ÄKTA pure SN2626712 (Cytiva) and UNICORN v7.11 (Cytiva) for FPLC purification of histone proteins.

UCSF ChimeraX v1.9 (Meng et al., 2023) for structure visualization.

LI-COR Odyssey M, LI-COR Acquisition v1.2, and ImageJ v1.54f (Schneider et al., 2012) for western blot and gel imaging, data analysis, and visualization.

QuantStudio™ 5 (Thermo Fisher, Cat# A28574) and QuantStudio™ Design & Analysis v2.7 for QC of sequencing samples.

Inkscape v1.3.2 for figure generation.

## DATA AVAILABILITY

Sequencing data can be accessed at GEO accension number GSE295057.

Raw data and analysis code can be accessed at Zenodo DOI: 10.5281/zenodo.15360677.

Cryo-EM maps and atomic models can be accessed with PDB IDs: 9OGS, 9OGZ (UM Sat2R-P); 9OGR, 9OH0 (Me Sat2R-P); 9OH1 (UM 601L); 9OH2 (Me 601L) and EMD IDs: EMD-70474 (UM Sat2R-P); EMD-70473 (Me Sat2R-P); EMD-70480 (UM 601L); EMD-70481 (Me 601L).

## ACKNOWLEDGEMENTS

We thank Joanna Yeung, Justin Rendleman, Viviana Risca, and the rest of the Risca lab, as well as Richard Koche, for their invaluable advice and sharing of reagents for the sequencing and bioinformatics analysis. We thank members of the Funabiki lab for their continual advice, particularly Yiming Niu for Cryo-EM analysis help and Isabel Wassing for help in generating the α-H2A.Z antibody. We thank Dr. David Shechter for the XTC-2 cell line and α-H2A.X-F serums, Shixin Liu and Aaron Straight for plasmids, and Tarun Kapoor for use of the Refeyn Two^MP^. This work was conducted using the High-Performance Computing Resource Center, the Evelyn Gruss Lipper Cryo-Electron Microscopy Resource Center, and the Proteomics Resource Center at the Rockefeller University (with the help of Henrik Molina and Soeren Heissel for proteomics analysis). Animal husbandry was supported by the staff at the Rockefeller Comparative Bioscience Center.

## AUTHOR CONTRIBUTIONS

R.M.S. and H.F. conceived and designed the study. R.M.S. and Y.A. designed and performed the Cryo-EM analysis. R.M.S. and H.A.K. designed physiological experiments. H.A.K. created the custom α-SRCAP antibody, prepared XTC-2 nuclei, and purified pG-Tn5. R.M.S. conducted all experiments and analysis. R.M.S. and H.F. prepared the manuscript with input from H.A.K. and Y.A.

