## Supplementary Figures for "IMPACTS OF DNA METHYLATION ON H2A.Z DEPOSITION AND NUCLEOSOME STABILITY"

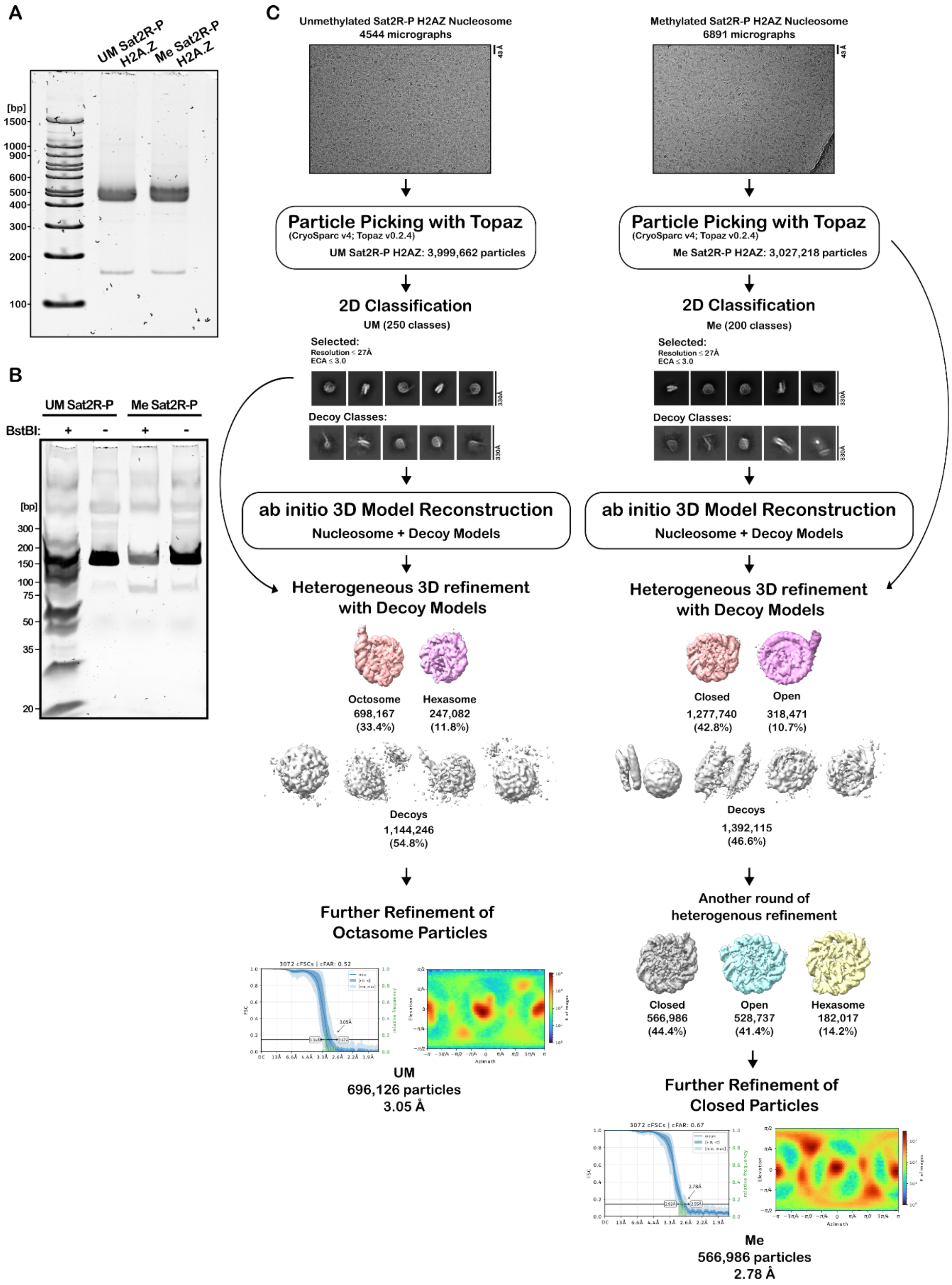


**Supplementary Figure 1. Workflow of Sat2R-P cryo-EM analysis. (A)** Native PAGE analysis of H2A.Z nucleosomes used for cryo-EM analysis. Bands visualized through SYBR Safe staining. **(B)** Native PAGE analysis of Sat2R-P DNA after digestion with BstBI to check for methylation status. **(C)** Diagram of structure analysis pipeline for either unmethylated (left) or methylated (right) H2A.Z SAT2R-P samples. Further details described in methods.

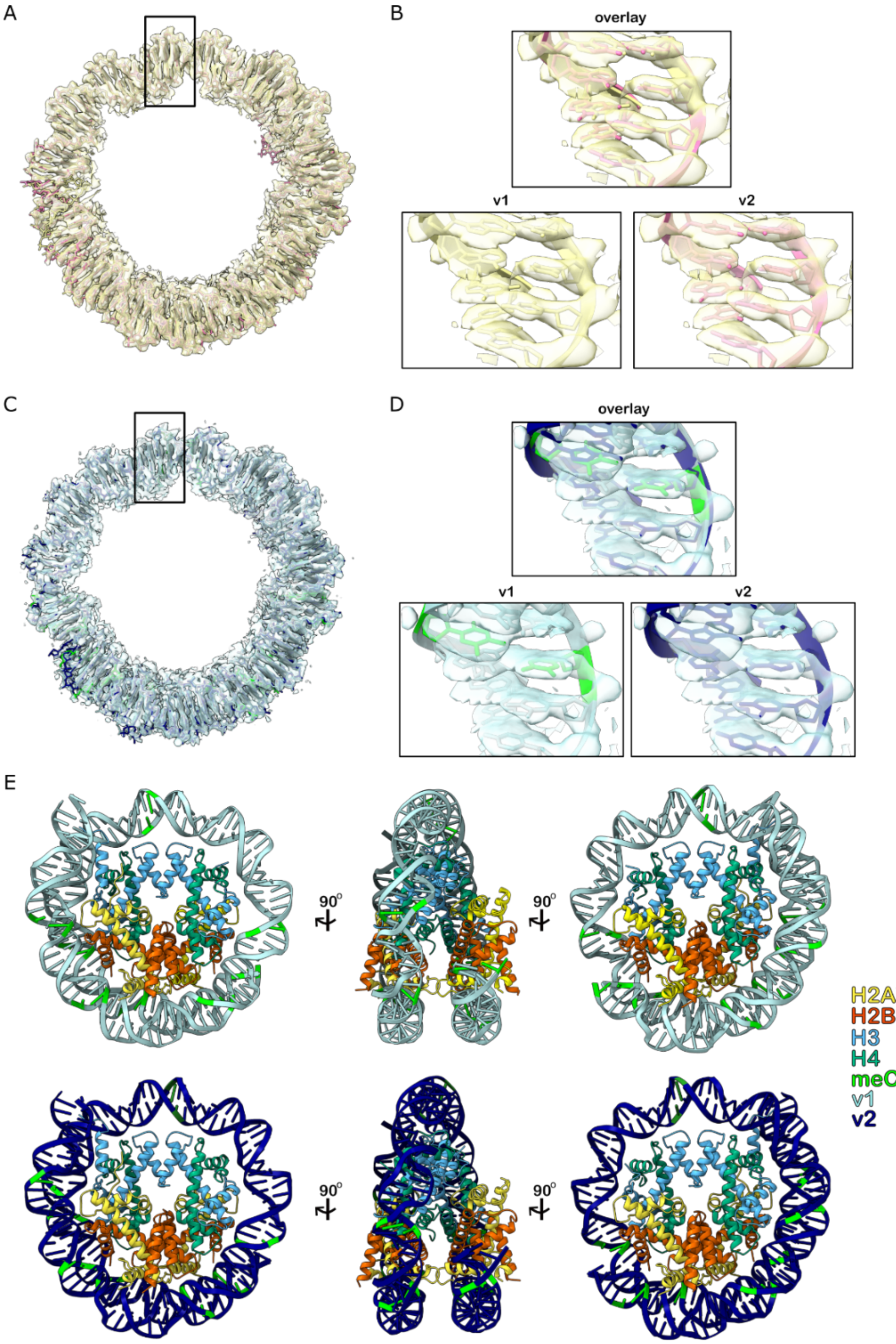


**Supplementary Figure 2. DNA atomic model generation for Sat2R-P H2A.Z nucleosome structures. (A and C)** Overlays of electron density maps for the DNA with both generated atomic models for either the unmethylated **(A)** or methylated **(C)** Sat2R-P H2A.Z nucleosome structure. **(B and D)** Zoom-in of dyad regions showing fitting of both DNA atomic models (v1 and v2) generated for either the unmethylated **(B)** or methylated **(D)** structure. Methylated cytosines are highlighted in green. Top panels are overlays of both models for the two structures. Positioning was determined by assessing densities around bases which retain purine/pyrimidine identity in both models vs those that switch. Bases that switched from purine to pyrimidine, and vice versa, showed more ambiguous densities than those that retained identity. Positioning was determined using the unmethylated structure and then used in modeling the methylated structure. (**E**) Comparison of predicted CpG positioning for the v1 (top) and v2 (bottom) atomic models generated for the methylated structure.

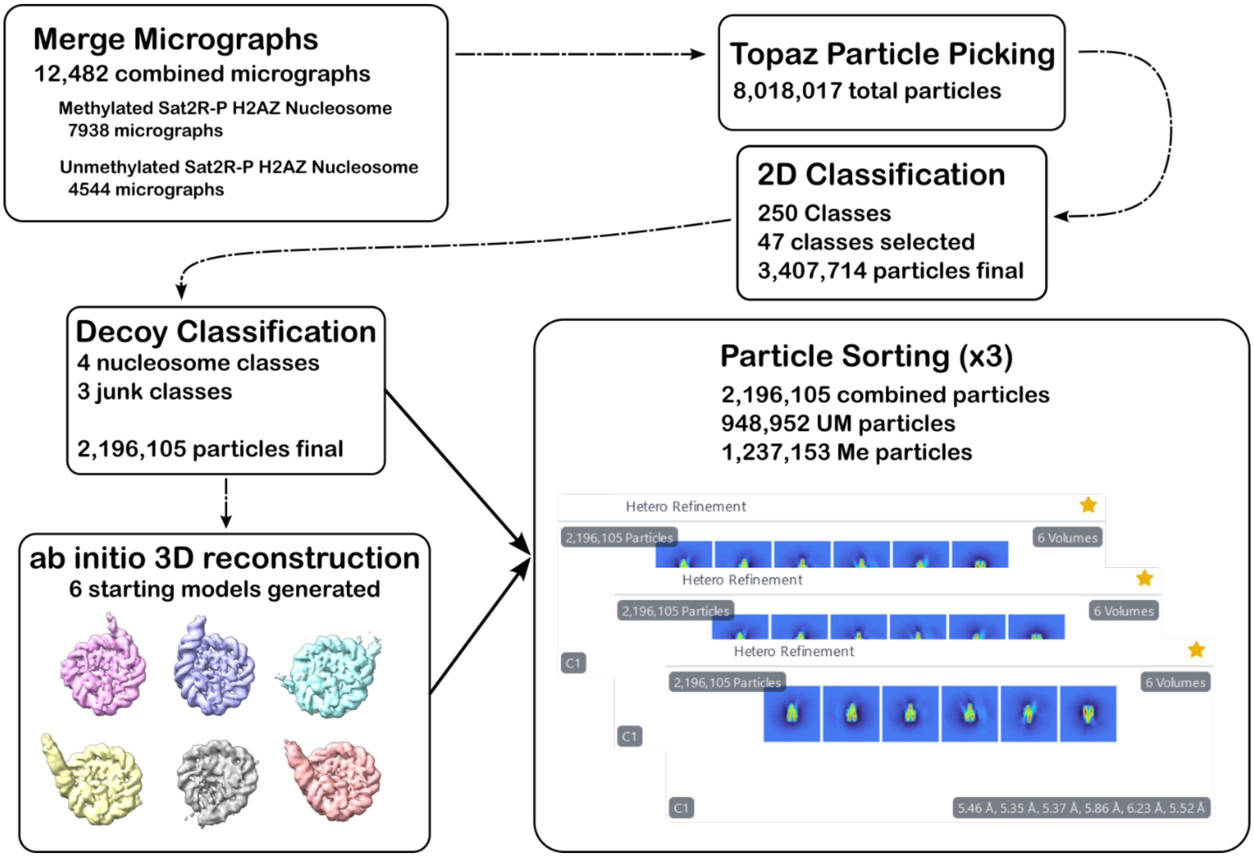


**Supplementary Figure 3. Workflow of *in silico* mixing 3D classification analysis.** Particles from a merged batch of unmethylated and methylated Sat2R-P H2A.Z nucleosome micrographs were picked and fed through the standard analysis pipeline. Six 3D models were then generated representing various open, closed, or hexasome-containing states of the nucleosome. Merged particles were then sorted to each of the classes through CryoSPARC's heterorefinement tool (n = 3 technical replicates).

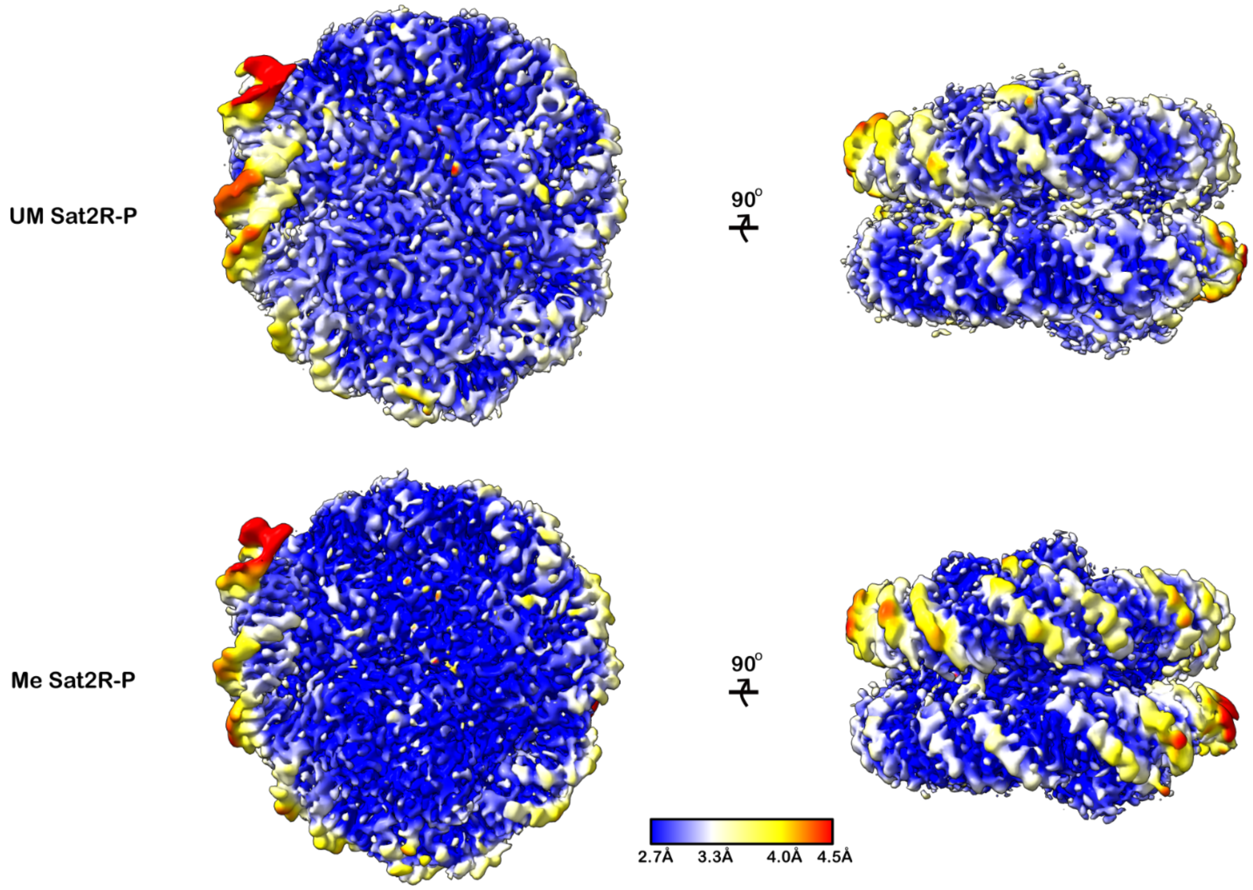


**Supplementary Figure 4. Local resolution estimates for unmethylated and methylated Sat2R-P H2A.Z structures.** Left, local resolution estimates of DF1 for both unmethylated (top) and methylated (bottom) Sat2R-P density maps. Right, side view showing local resolution of the DNA backbone for both methylated and unmethylated density maps. Local resolution maps were generated in CryoSPARC using the local resolution estimation tool and overlaid on density maps filtered using CryoSPARC's local filtering tool.

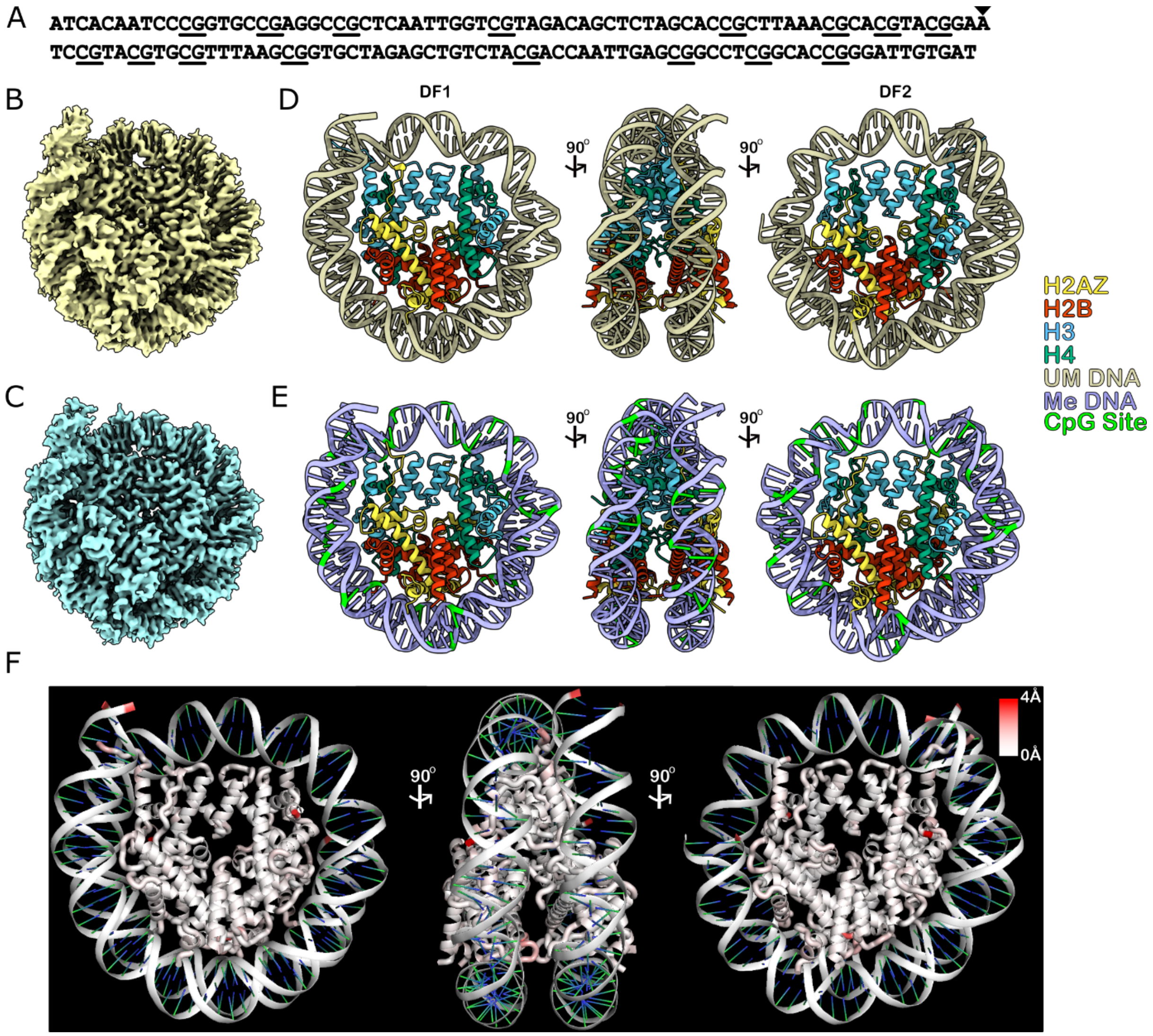


**Supplementary Figure 5. Cryo-EM structures of 601L H2A.Z nucleosomes show no DNA methylation dependent differences. (A)** Diagram of the 205 bp palindromic 601L sequence. Triangle indicates sequence midpoint. CpGs are underlined. **(B and C)** Final density maps of unmethylated **(B)** or methylated **(C)** H2A.Z nucleosomes. **(D and E)** Atomic models generated for **(B and C)**. Methylated cytosines shown in green **(F)** RMSD analysis comparing differences between methylated and unmethylated 601L H2A.Z atomic models.

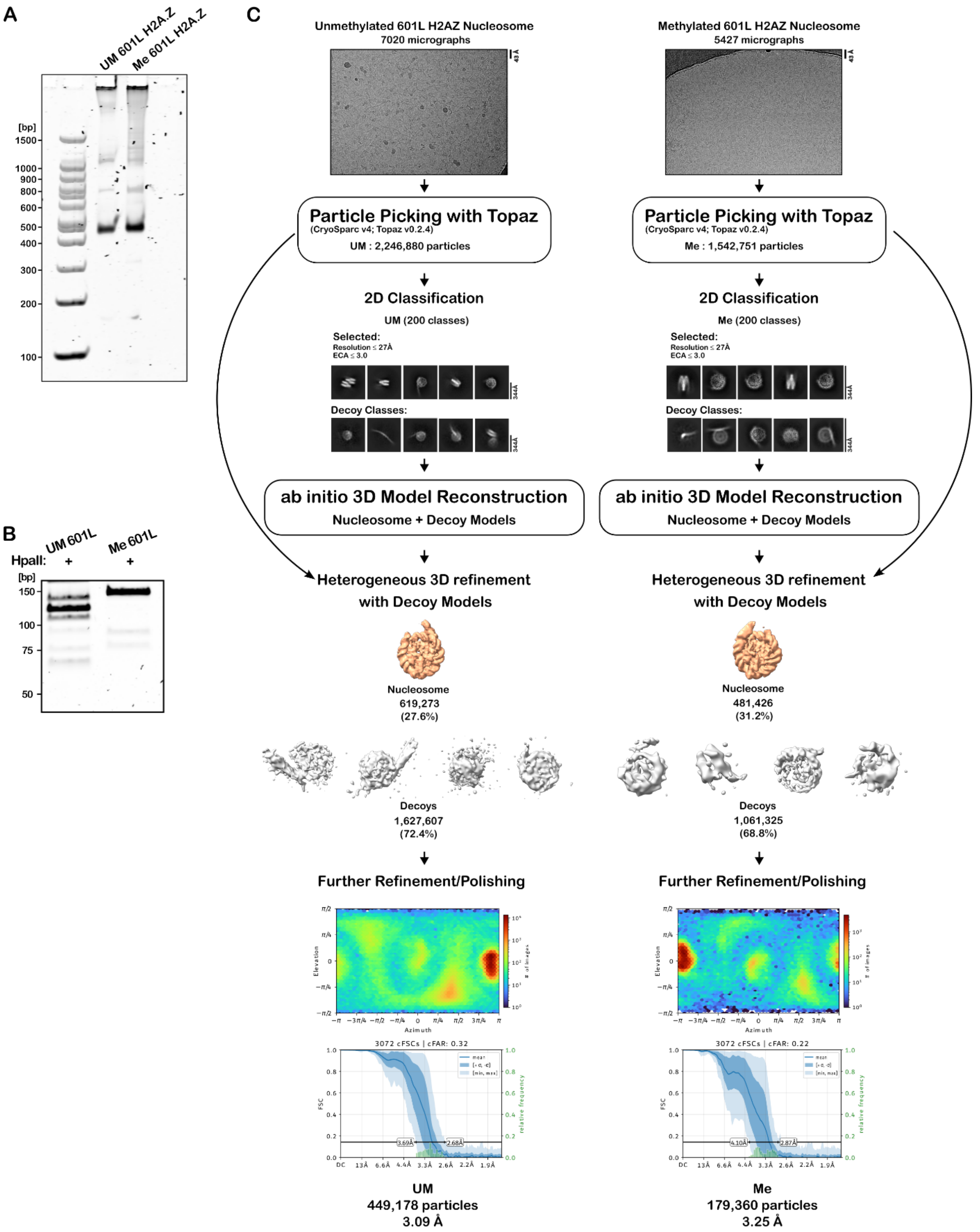


**Supplementary Figure 6. Workflow of 601L H2A.Z nucleosome cryo-EM structures. (A)** Native PAGE analysis of 601L H2A.Z nucleosome samples used for cryo-EM. Bands visualized through SYBR Safe staining. **(B)** Native PAGE analysis of 601L DNA after digestion with HpaII to check for methylation status. **(C)** Diagram showing analysis pipeline for solving 601L nucleosome structures. Further details described in methods.

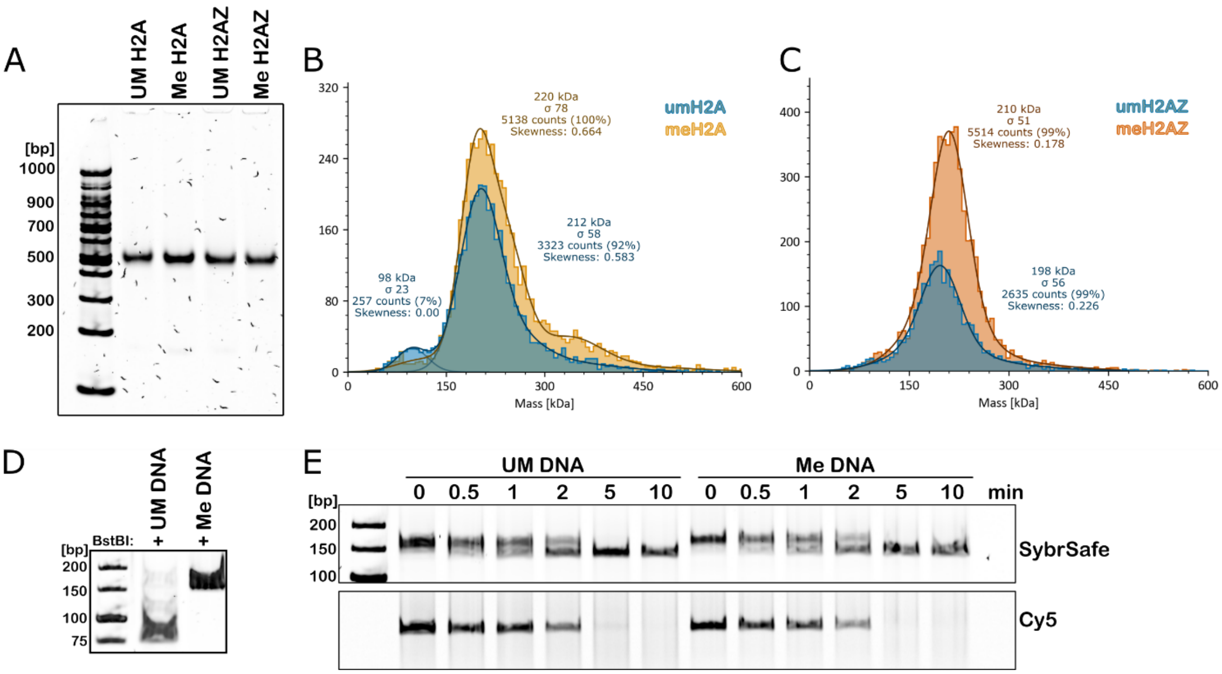


**Supplementary Figure 7. Native PAGE and mass photometry analysis of nucleosomes used in HinfI digest assays. (A)** Native PAGE analysis of nucleosome samples. Bands visualized through SYBR Safe staining. **(B and C)** Mass photometry analysis of methylated or unmethylated H2A **(B)** or H2A.Z **(C)** nucleosomes. Peaks around 200-220 kDa represent the fully formed nucleosome population. **(D)** BstBI digestion of 1HinfI_Sat2R DNA to verify methylation status. **(E)** HinfI digest time course of bare unmethylated or methylated DNA.

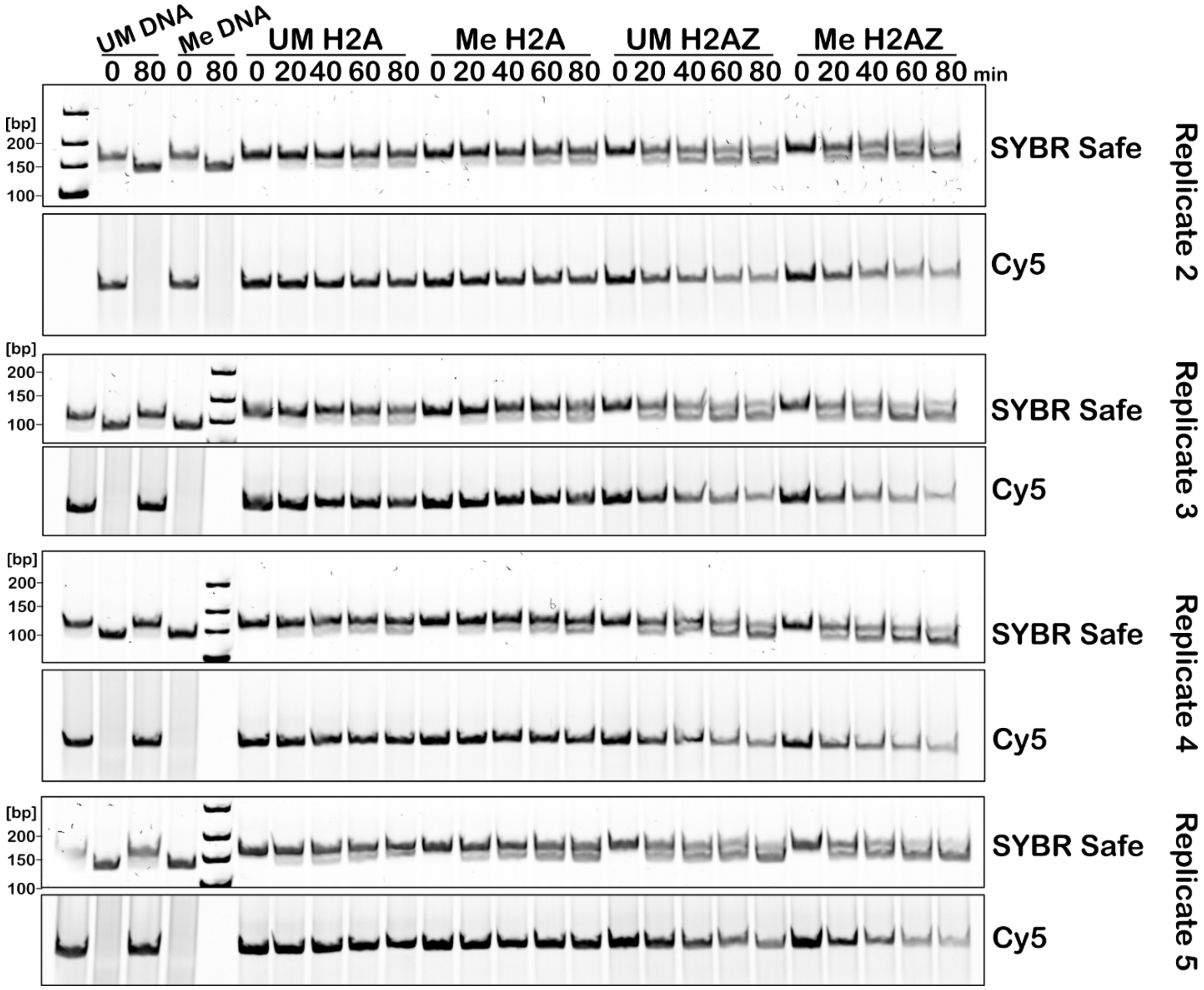


**Supplementary Figure 8. Replicate gels of data from Main Figure 3.**

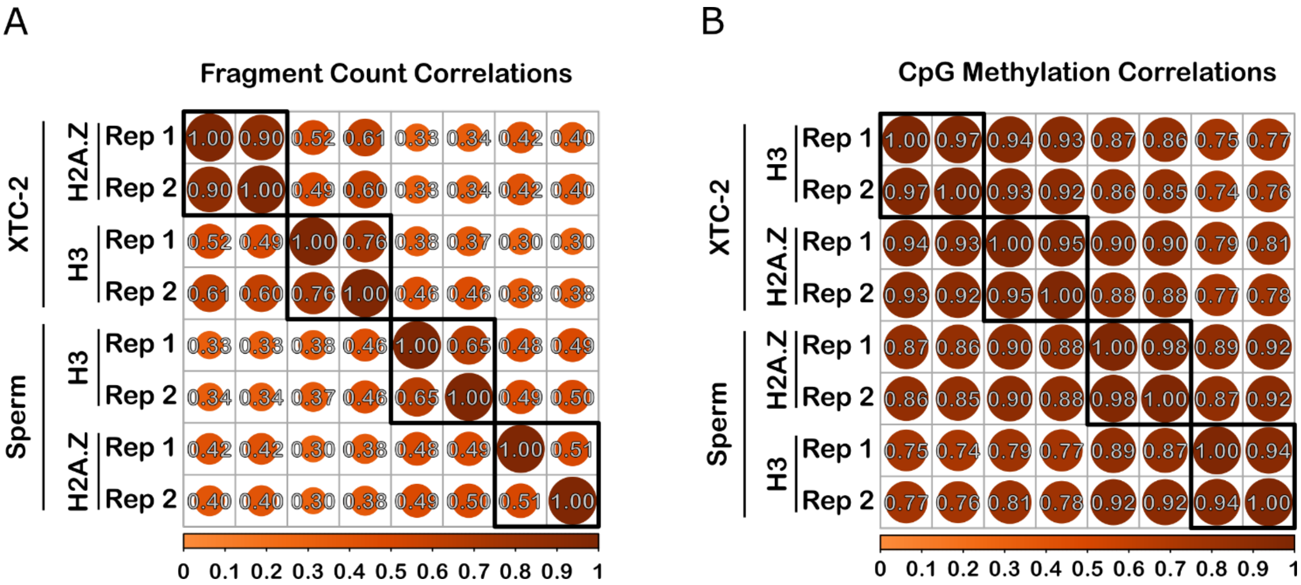


**Supplementary Figure 9. Reproducibility comparison of CnT-BS samples (A)** Pearson correlation plot comparing fragment counts mapped to 1000 bp genomic bins across samples. Bins containing zero fragments were first filtered out. Fragment counts were then normalized using DESeq2 to account for sequencing depth differences and underwent a variance stabilizing transformation (VST) before the correlation matrix was generated. **(B)** Pearson correlation plot comparing CpG methylation across samples. Percent methylation at individual CpG sites was calculated for high coverage CpGs (≥ 10 reads) and normalized using methylKit. The correlation matrix was then generated using pairwise observations across all eight samples. Both correlation matrices were ordered, and rectangles drawn (n = 4, user-specified), via hierarchical clustering.

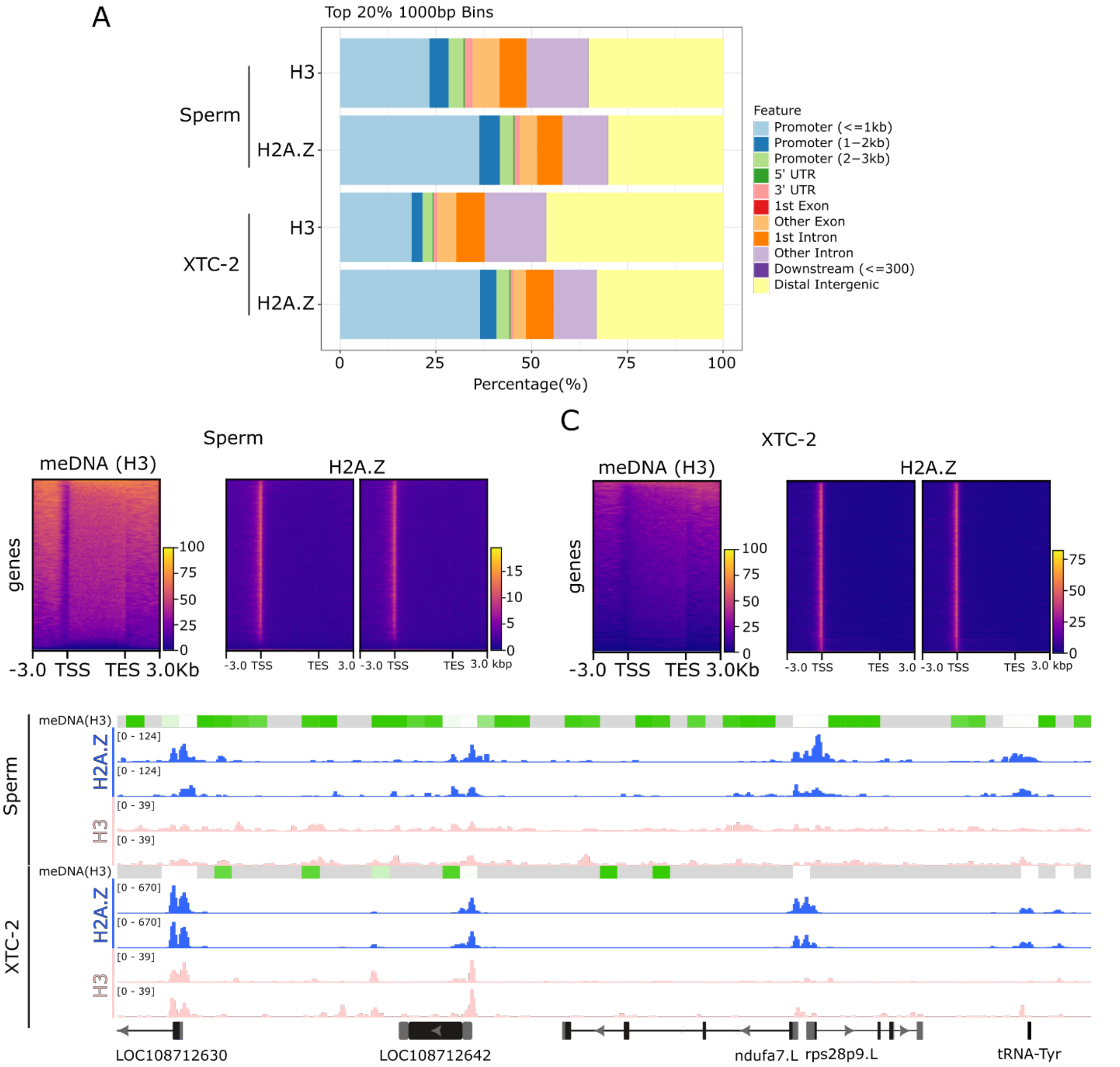


**Supplementary Figure 10. H2A.Z localizes to hypomethylated TSSs. (A)** Genomic annotation plots for the top quantile of 1000 bp genomic bins containing the most mapped reads averaged between replicates for each specified condition. **(B and C)** Heatmaps of either H3 CpG methylation calls averaged over 500 bp tiles (left-most panel) or H2A.Z reads (right panels) from two biological replicates across a representation of all annotated *Xenopus laevis* genes from either sperm (**B**) or XTC-2 (**C**) samples. Corresponding H3 and H2A.Z data are shown on the same sets of genes sorted in the same order. **(D)** IGV snapshot of a region on chromosome 1L showcasing preferential H2A.Z deposition at hypomethylated TSSs. Average H3 methylation tracks were generated across 500 bp tiles for visualization purposes on a scale of 0 to 100% CpG methylation. Grey indicates regions with no methylation data availability. Two biological replicates of H2A.Z and H3 tracks are shown.

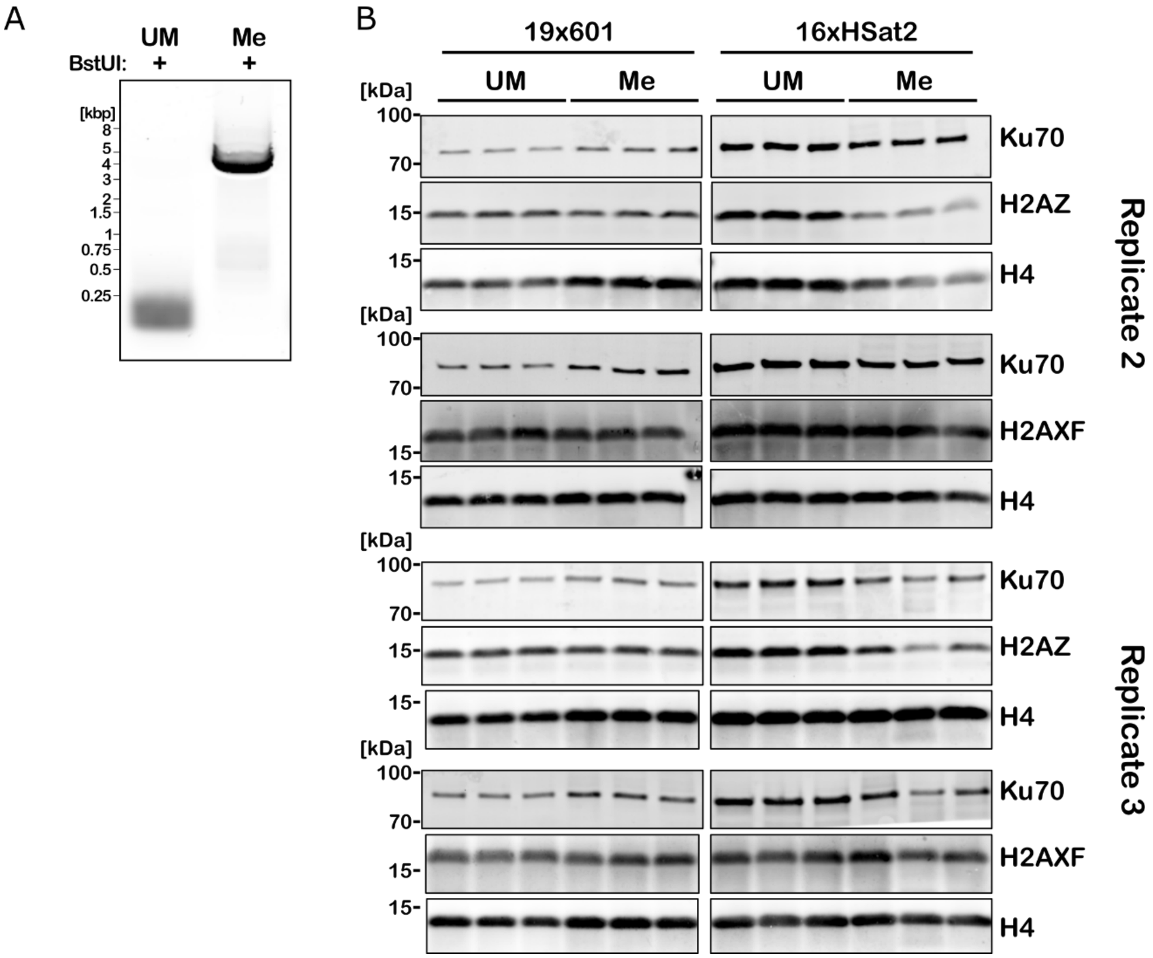


**Supplementary Figure 11. DNA methylation check and replicates of data from Main Figure 4. (A)** BstUI digestion of 19x601 DNA substrates to verify methylation status. Both 19x601 and 16xHSat2 DNA were methylated under identical conditions, however the HSat2 sequence lacks appropriate cut sites for methylation-sensitive restriction enzymes. The 601 substrates were used to determine complete methylation status for each batch. **(B)** Replicate blots for data shown in **Main Figure 4**.

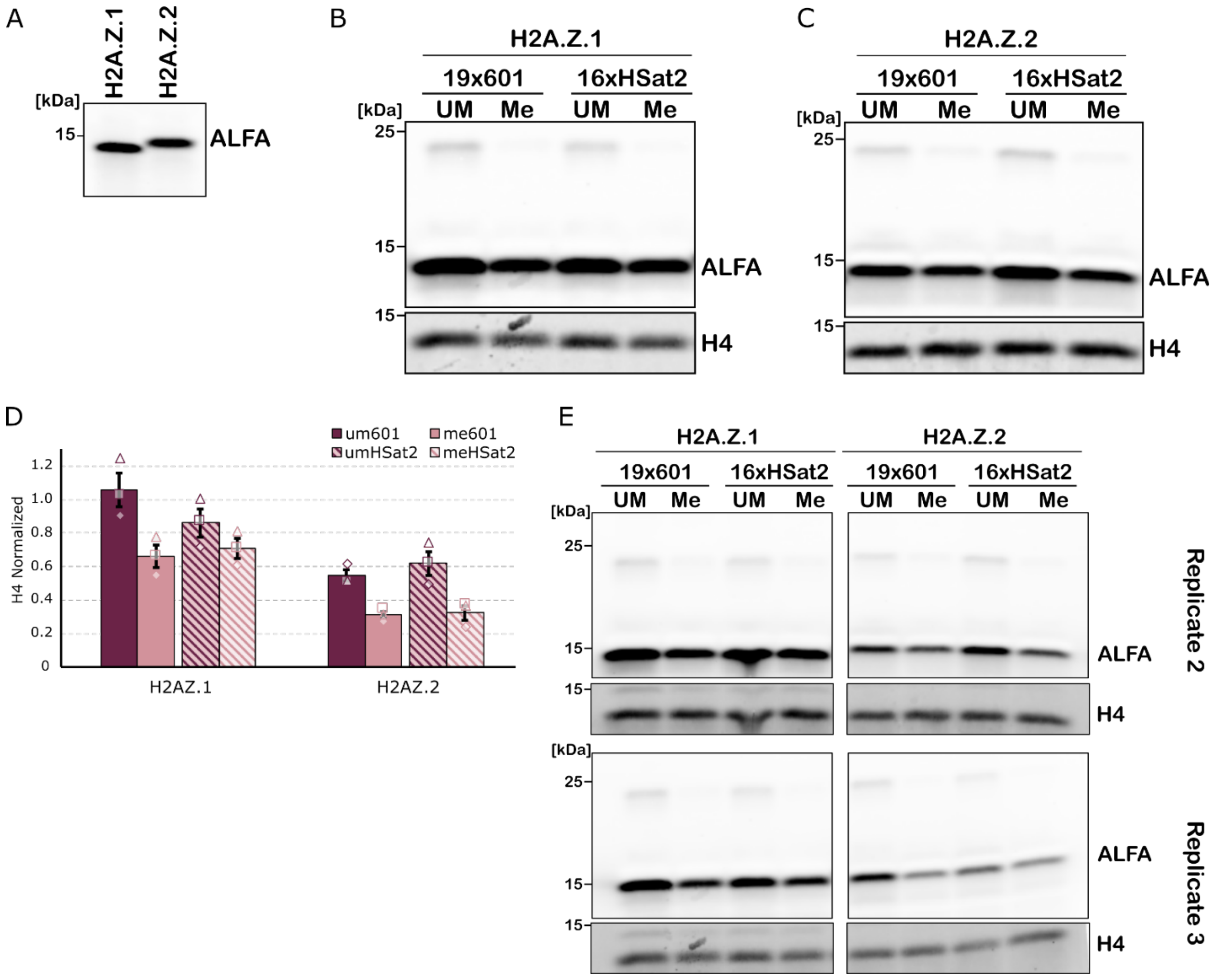


**Supplementary Figure 12. Both H2A.Z variants preferentially associate with unmethylated DNA substrates. (A)** Expression check of exogenously added mRNA expression for ALFA-tagged versions of both H2A.Z.1 and H2A.Z.2. **(B and C)** Blots of DNA beads incubated with interphase extract expressing either ALFA-tagged H2A.Z.1 **(B)** and H2A.Z.2 **(C).** Presence of the specified H2A.Z variant on DNA beads was determined by blotting against the ALFA-tag with H4 shown as a loading control. **(D)** Quantification of **(B and C)**. ALFA signals were normalized to H4 signals. Unique shapes denote results from three independent experiments (n = 3 biological replicates). Error bars represent SEM. **(E)** Additional replicate blots of data from **(D)**.

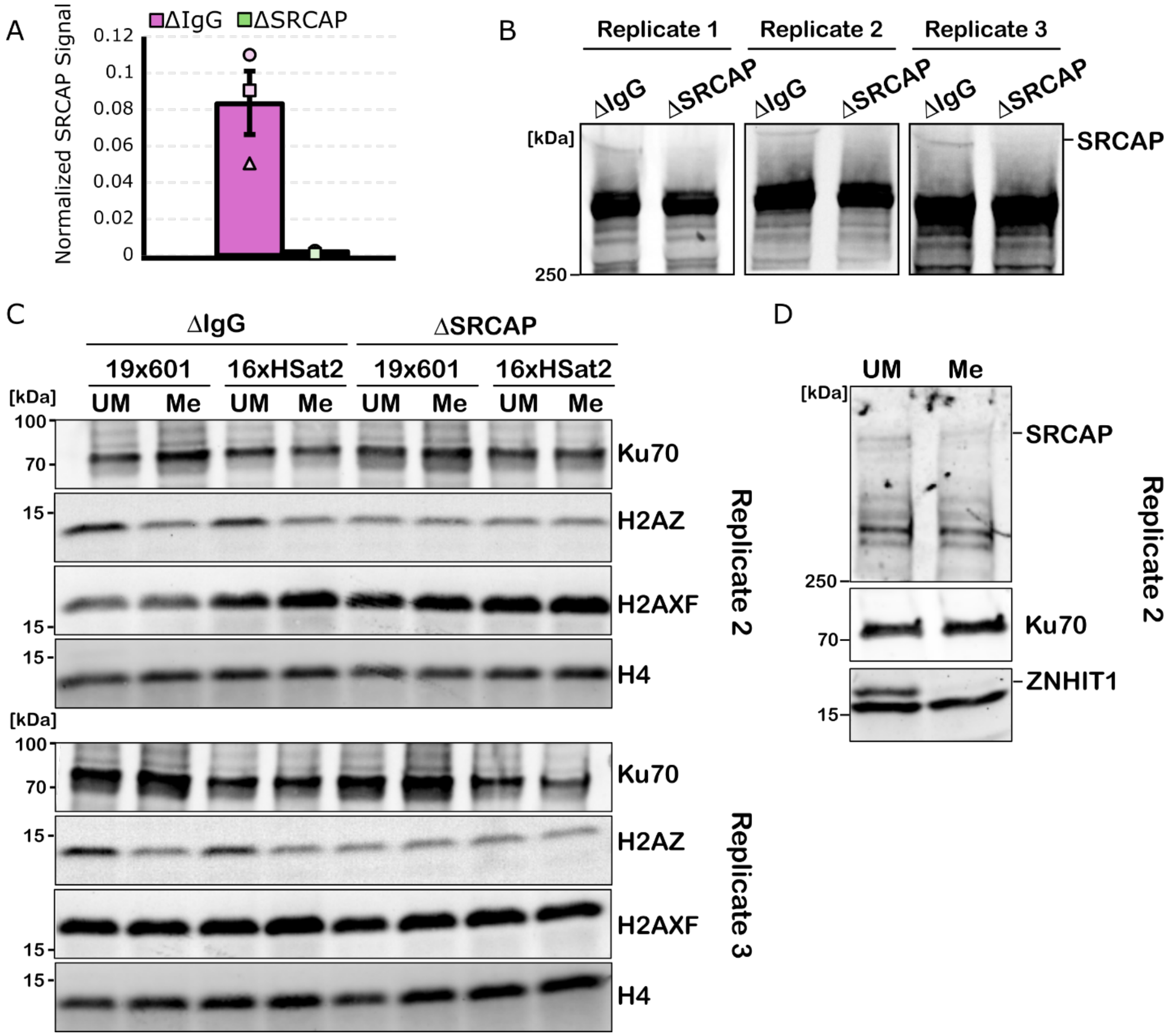


**Supplementary Figure 13. SRCAP depletion efficiency and replicates of data from Main Figure 5. (A)** Quantification of SRCAP WB signal from egg extract after depletion with anti-IgG or anti-SRCAP antibody beads. Replicates from three independent experiments shown. Error bars represent SEM. **(B)** Replicate blots of data plotted in (**A**). **(C)** Replicate blots of **Main Figure 5B-D**. **(D)** Replicate blots of **Main Figure 5E-F** with Ku70 as a loading control.

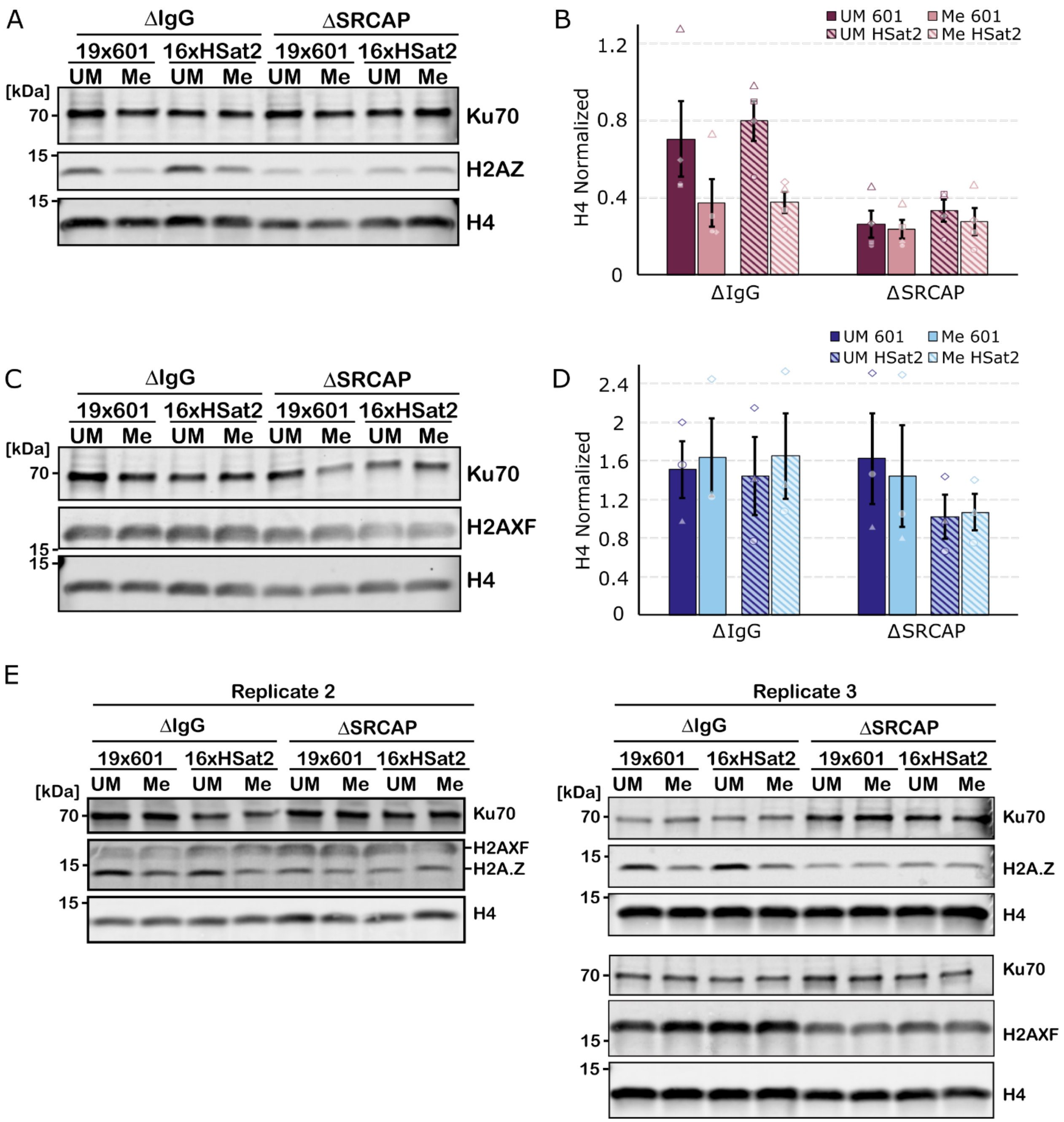


**Supplementary Figure 14. Chromatinization assay with SRCAP depletion using a commercial antibody. (A and C)** Western blots probing for H2A.Z (**A**) or H2A.X-F (**C**) signal along with respective loading controls bound to DNA beads incubated in interphase egg extract after SRCAP depletion using a commercial antibody raised against human SRCAP (Kerafast, Cat#: ESL103). **(B and D)** Quantification of (**A** and **C**). Results from three independent experiments plotted. Error bars represent SEM. **(E)** Replicate western blots.

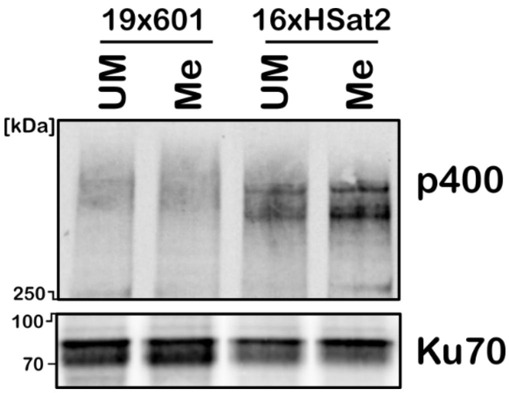


**Supplementary Figure 15. TIP60-C does not display DNA methylation sensitivity.** Western blots of DNA bead pulldowns from interphase egg extract probing for endogenous p400 (ATPase for Tip60-C) along with respective loading controls.

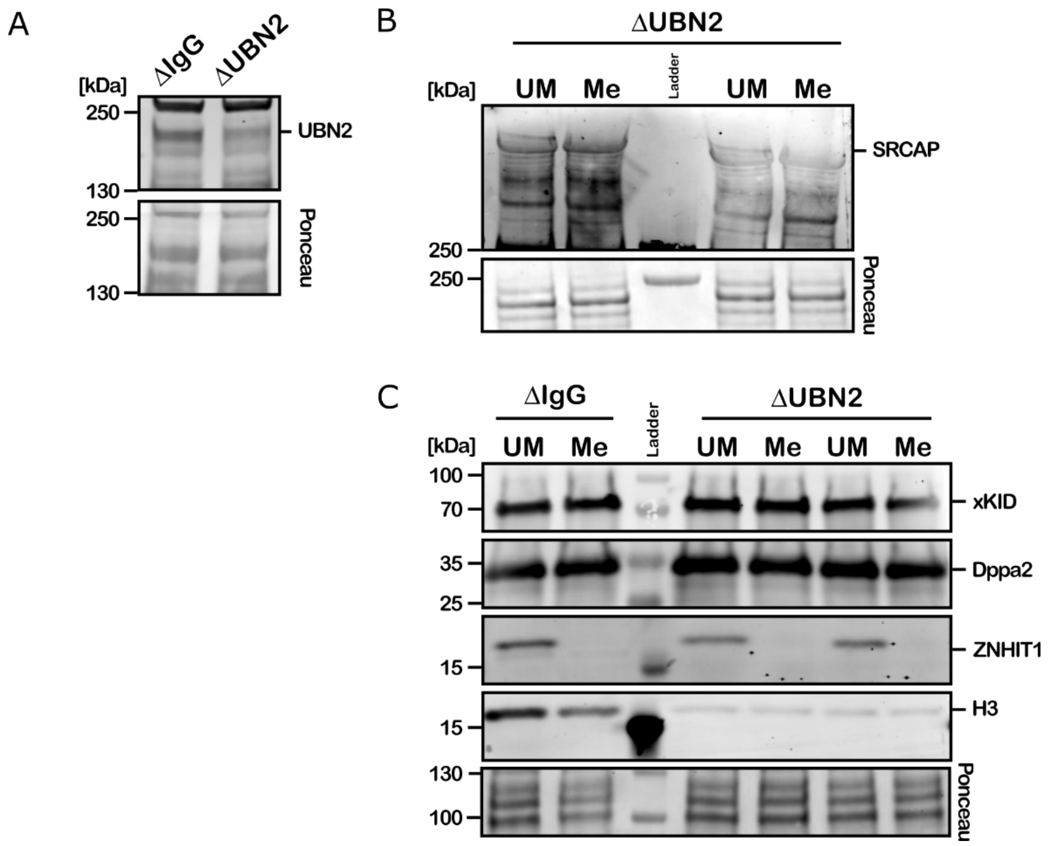


**Supplementary Figure 16. SRCAP-C maintains unmethylated DNA binding bias in the absence of nucleosome loading. (A)** Western blot showing UBN2 depletion efficiency. **(B)** Western blot of SRCAP on DNA beads incubated in UBN2-depleted extract. Two replicates shown and total protein via ponceau staining used to assess loading. **(C)** Western blot of DNA beads incubated in IgG control-depleted and UBN2-depleted extract. ZNHIT1, subunit of SRCAP-C, displays preference for unmethylated DNA across conditions. DNA binding proteins Dppa2 and xKID did not show any differences in chromatin accessibility between DNA methylation status. H3 shown to verify decreased nucleosome loading upon UBN2 depletion. Total protein via ponceau staining used to assess loading. One replicate of ΔIgG and two replicates of ΔUBN2 conditions shown.
