## Supplementary Tables for "IMPACTS OF DNA METHYLATION ON H2A.Z DEPOSITION AND NUCLEOSOME STABILITY"

**Supplementary Table 1. Collection and model statistics for the Cryo-EM structures.**

| **Sample** | **UM Sat3R-P** | **Me Sat2R-P** | **UM 601L** | **Me 601L** |
|---|---|---|---|---|
| **Data Collection** | | | | |
| Microscope | Titan Krios | | | |
| Magnification | 81,000 | | | |
| Voltage (kV) | 300 | | | |
| Camera | Gatan K3 | | | |
| Electron exposure (e-/Å) | 50.703 | | 50 | |
| Defocus range (µm) | -1.5 ~ -3.5 | | | |
| Pixel size (Å) | 0.86 | | | |
| Micrographs Collected | 4544 | 6891 | 7020 | 5427 |
| **Data Processing** | | | | |
| Particles (no.) | 696,126 | 566,986 | 449,178 | 179,360 |
| Symmetry imposed | C1 | C1 | C1 | C1 |
| Map resolution (Å, FSC = 0.143) | 3.05 | 2.78 | 3.09 | 3.25 |
| ***EMDR ID*** | ***EMD-70474*** | ***EMD-70473*** | ***EMD-70480*** | ***EMD-70481*** |

| | **UM Sat2R-P** | | **Me Sat2R-P** | | **601L** | |
|---|---|---|---|---|---|---|
| **Atomic Models** | **v1** | **v2** | **v1** | **v2** | **UM** | **Me** |
| *Composition* | | | | | | |
| Chains | 10 | 10 | 10 | 10 | 10 | 10 |
| Atoms | 11117 (Hydrogens: 0) | 10905 (Hydrogens: 0) | 10585 (Hydrogens: 0) | 10558 (Hydrogens: 0) | 11101 (Hydrogens: 0) | 11477 (Hydrogens: 0) |
| Residues | Protein: 735 Nucleotide: 259 | Protein: 721 Nucleotide: 254 | Protein: 701 Nucleotide: 246 | Protein: 698 Nucleotide: 246 | Protein: 722 Nucleotide: 264 | Protein: 738 Nucleotide: 274 |
| Water | 0 | 0 | 0 | 0 | 0 | 0 |
| Ligands | 0 | 0 | 0 | 0 | 0 | 0 |
| Bonds (RMSD) | | | | | | |
| Length (Å) (# > 4σ) | 0.004 (0) | 0.003 (0) | 0.004 (0) | 0.003 (0) | 0.003 (0) | 0.004 (0) |
| Angles (°) (# > 4σ) | 0.580 (0) | 0.525 (3) | 0.617 (0) | 0.619 (0) | 0.578 (1) | 0.650 (0) |
| MolProbity score | 1.78 | 1.47 | 1.77 | 1.82 | 1.6 | 2.35 |
| Clash score | 5.17 | 4.66 | 5.86 | 7.13 | 8.64 | 9.79 |
| Ramachandran plot (%) | | | | | | |
| Outliers | 0 | 0 | 0 | 0 | 0.14 | 0 |
| Allowed | 2.5 | 1.56 | 1.9 | 2.35 | 1.56 | 4.16 |
| Favored | 97.5 | 98.44 | 98.1 | 97.65 | 98.3 | 95.84 |
| Rama-Z (Ramachandran plot Z-score RMSD) | | | | | | |
| whole (N = 719) | 1.58 (0.32) | 1.80 (0.32) | 1.67 (0.32) | 1.84 (0.32) | 1.99 (0.32) | 0.95 (0.30) |
| helix (N = 518) | 2.25 (0.23) | 2.33 (0.23) | 2.07 (0.23) | 2.20 (0.23) | 2.25 (0.22) | 1.63 (0.22) |
| sheet (N = 0) | --- (---) | --- (---) | --- (---) | --- (---) | --- (---) | --- (---) |
| loop (N = 201) | -2.34 (0.36) | -2.21 (0.36) | -2.05 (0.39) | -1.87 (0.38) | -1.43 (0.42) | -2.12 (0.36) |
| Rotamer outliers (%) | 3.44 | 1.99 | 3.96 | 2.94 | 1.5 | 5.37 |
| Cβ outliers (%) | NA | NA | NA | NA | NA | NA |
| Peptide plane (%) | | | | | | |
| Cis proline/general | 0.0/0.0 | 0.0/0.0 | 0.0/0.0 | 0.0/0.0 | 0.0/0.0 | 0.0/0.0 |
| Twisted proline/general | 0.0/0.0 | 0.0/0.0 | 0.0/0.0 | 0.0/0.0 | 0.0/0.0 | 0.0/0.0 |
| CaBLAM outliers (%) | 1.85 | 1.74 | 0.9 | 1.2 | 1.01 | 1.42 |
| ADP (B-factors) | | | | | | |
| Iso/Aniso (#) | 11117/0 | 10905/0 | 10585/0 | 10557/0 | 11103/0 | 11477/0 |
| min/max/mean | | | | | | |

| | | | | | | |
|---|---|---|---|---|---|---|
| Protein | 0.00/41.12/5.00 | 0.28/93.40/15.80 | 0.23/30.35/4.02 | 0/89.15/13.38 | 1.75/82.28/22.40 | 2.38/89.91/24.49 |
| Nucleotide | 0.23/83.07/30.56 | 3.71/150.32/64.25 | -0.00/88.15/40.33 | 0.06/161.03/73.80 | 4.85/140.55/59.47 | 6.08/156.41/67.10 |
| Ligand | --- | --- | --- | --- | --- | --- |
| Water | --- | --- | --- | --- | --- | --- |
| Occupancy | | | | | | |
| Mean | 1 | 1 | 1 | 1 | 1 | 1 |
| occ = 1 (%) | 100 | 100 | 100 | 100 | 100 | 100 |
| 0 < occ < 1 (%) | 0 | 0 | 0 | 0 | 0 | 0 |
| occ > 1 (%) | 0 | 0 | 0 | 0 | 0 | 0 |
| *Box* | | | | | | |
| Lengths (Å) | 79.12, 119.54, 119.54 | 78.26, 118.68, 118.68 | 77.4, 116.96, 117.82 | 77.4, 116.96, 117.82 | 75.68, 116.1, 122.98 | 73.96, 116.1, 121.26 |
| Angles (°) | 90, 90, 90 | 90, 90, 90 | 90, 90, 90 | 90, 90, 90 | 90, 90, 90 | 90, 90, 90 |
| Supplied Resolution (Å) | 3.3 | 3.3 | 3 | 3 | 3.2 | 3.3 |
| Resolution Estimates (Å) | Masked \| Unmasked | Masked \| Unmasked | Masked \| Unmasked | Masked \| Unmasked | Masked \| Unmasked | Masked \| Unmasked |
| d FSC (half maps; 0.143) | --- \| --- | --- \| --- | --- \| --- | --- \| --- | --- \| --- | --- \| --- |
| d 99 (full/half1/half2) | 3.5/---/--- \| 3.5/---/--- | 3.5/---/--- \| 3.5/---/--- | 3.2/---/--- \| 3.2/---/--- | 3.2/---/--- \| 3.2/---/--- | 3.3/---/--- \| 3.3/---/--- | 3.4/---/--- \| 3.3/---/--- |
| d model | 3.4 \| 3.4 | 3.4 \| 3.4 | 3.1 \| 3.2 | 3.1 \| 3.2 | 3.2 \| 3.2 | 3.3 \| 3.3 |
| d FSC model (0/0.143/0.5) | 2.9/3.1/3.3 \| 3.1/3.2/3.3 | 2.9/3.1/3.3 \| 3.0/3.2/3.3 | 2.6/2.9/3.0 \| 2.8/3.0/3.1 | 2.6/2.9/3.0 \| 2.8/3.0/3.1 | 2.9/3.0/3.2 \| 2.9/3.1/3.3 | 3.1/3.1/3.3 \| 3.1/3.2/3.4 |
| Map min/max/mean | -39.16/62.30/0.79 | -39.16/62.3/0.82 | -41.24/58.89/0.40 | -41.24/58.89/0.40 | -34.81/61.05/0.68 | -31.70/51.07/0.56 |
| *Model vs Data* | | | | | | |
| CC (mask) | 0.86 | 0.88 | 0.83 | 0.83 | 0.82 | 0.8 |
| CC (box) | 0.78 | 0.78 | 0.7 | 0.68 | 0.74 | 0.7 |
| CC (peaks) | 0.75 | 0.75 | 0.68 | 0.66 | 0.7 | 0.65 |
| CC (volume) | 0.8 | 0.82 | 0.75 | 0.76 | 0.78 | 0.76 |
| Mean CC for ligands | --- | --- | --- | --- | --- | --- |
| ***PDB ID*** | ***9OGS*** | ***9OGZ*** | ***9OGR*** | ***9OH0*** | ***9OH1*** | ***9OH2*** |

**Supplementary Table 2. Alignment statistics of CnT-BS sequencing libraries.** XTC reads were downsampled to 30 million reads to match sperm samples, then all raw reads were trimmed and deduplicated using fastp v0.24.0 and aligned to the *Xenopus* laevis genome (Xenla 10.1) or lambda genome (to assess for bisulfite conversion efficiency) using Bismark v0.24.2. Methylated CpG percentages for H2A.Z samples were determined for the top 2.5 % of H2A.Z peaks by AUC (obtained using SEACR v1.3) while methylated percentages for H3 associated CpGs was conducted on regions outside of determined H2A.Z peaks. All CpGs met a cutoff of at least 5 reads to be counted. Methylated lambda CpG statistics were calculated from all mapped CpGs with no filtering.

| | | *Alignment to Xenopus laevis* | | | *Alignment to lambda* | |
|---|---|---|---|---|---|---|
| | | **Aligned %** | **Total Aligned Reads (No.)** | **Methylated CpG %** | **Aligned %** | **Methylated CpG %** |
| **Sperm Pronuclei** | | | | | | |
| **H3** | *Rep 1* | 71.7 | 24141516 | **85.6** | 0.8 | **0.6** |
| | *Rep 2* | 71.0 | 24185147 | **85** | 1.0 | **0.6** |
| **H2AZ** | *Rep 1* | 64.7 | 25362485 | **42.5** | 4.0 | **0.6** |
| | *Rep 2* | 69.0 | 21963635 | **43.5** | 2.5 | **0.6** |
| **XTC-2** | | | | | | |
| **H3** | *Rep 1* | 68.2 | 19539297 | **53.2** | 7.2 | **0.4** |
| | *Rep 2* | 62.4 | 17758471 | **53.8** | 15.9 | **0.3** |
| **H2AZ** | *Rep 1* | 69.1 | 19409396 | **3.2** | 12.4 | **0.3** |
| | *Rep 2* | 70.9 | 20109208 | **2.8** | 9.8 | **0.3** |

**Supplementary Table 3. Fragment statistics of filtered genomic bins used for sequencing analysis.** A count matrix was generated from processed bam files over 1,000 bp bins of the *Xenopus laevis* genome. Mitochondrial reads and bins containing fewer than 17 reads across less than 2 samples were removed to filter out low signal areas. Replicates for each sample were averaged and resulting count matrix used for genomic annotation analysis.

| | | Fragments in Bins (No.) | Fragments in Bins (%) |
|---|---|---|---|
| **Sperm Pronuclei** | | | |
| **H3** | *Rep 1* | 9062125 | 34.04 |
| | *Rep 2* | 9265292 | 34.68 |
| **H2AZ** | *Rep 1* | 11482371 | 43.04 |
| | *Rep 2* | 9213698 | 39.58 |
| **XTC-2** | | | |
| **H3** | *Rep 1* | 7789646 | 36.43 |
| | *Rep 2* | 8145900 | 41.79 |
| **H2AZ** | *Rep 1* | 14276029 | 66.62 |
| | *Rep 2* | 15187573 | 69.13 |
| **Total Bins (No.)** | | 506009 | |
